# Partition Coefficients Reveal Changes in Properties of Low-Contrast Biomolecular Condensates

**DOI:** 10.64898/2026.02.20.707107

**Authors:** Kaarthik Varma, Dana Matthias, Chase B. Shapiro, Sullivan Bailey-Darland, Takumi Matsuzawa, Charlotta Lorenz, Teagan Bate, Stephen J. Thornton, Senthilkumar Duraivel, Robert W. Style, James P. Sethna, Eric R. Dufresne

## Abstract

Biomolecular condensates are domains within cells with distinct compositions, held together by intermolecular cohesion. They are implicated in a variety of cellular processes, and *in vitro* studies have revealed the molecular driving forces that underly their condensation. However, *in vitro* condensates do not capture essential features of cellular condensates. In particular, enrichment of proteins, quantified by partition coefficients, is often exaggerated in these simplified systems. We show that the addition of free amino acids and other small molecules to model condensates can bring their partition coefficients within physiological range. In this limit, where there is low biochemical contrast between condensates and their surroundings, we observe striking changes to condensate behavior. Such low-contrast condensates exhibit large fluctuations in shape and composition and show enhanced sensitivity to changes in their environment. These behaviors reflect dramatic shifts to their material properties, including interfacial tension, rheology, and chemical susceptibilities. We note remarkable similarities in these effects across seemingly unrelated two-phase fluid systems. To explain these trends, we reformulate classic models of critical phenomena in terms of partition coefficients. This framework simplifies application of theory to experiments with near-critical fluids and suggests new experimental approaches for assessing condensate physiology in live cells.

## I. INTRODUCTION

Biomolecular condensates organize the cytoplasm into distinct domains [1] whose chemical micro-evironments and material properties enable diverse physiology [2–7]. While studies of simplified systems with a few molecular components have revealed sequence-specific interaction motifs that drive condensation [8–13], physiological phase separation occurs in a different context, featuring competing interactions from thousands of different proteins, nucleic acids, and metabolites [14–17]. These molecular by-standers can have pronounced effects on the stability and material properties of condensates [18–25], confounding the link between *in vitro* and *in vivo* experiments. Since adjusting *in vitro* conditions to exactly match the complex crowded environment of living cells would be a futile exercise, we need new experimental and conceptual frameworks to dissect condensate function.

Here, we reveal a direct link between condensate composition and material properties based on partition coefficients, which quantify the chemical contrast of condensates with their surroundings. Lowering the partition coefficients of diverse *in vitro* condensates into physiological range, we found marked changes to their behavior, including enhanced sensitivity to perturbations and dynamic fluctuations in shape and composition. Despite diverse condensation mechanisms, we found common trends in the underlying phase diagrams and material properties. These results can be understood using classic theories of low-contrast multi-phase materials. The unusual properties of condensates at low contrast suggest new mechanisms for their regulation and function in cells, and can be exploited to narrow the gap between *in vitro* and *in vivo* observations.

## RESULTS AND DISCUSSION

### A. Cellular condensates have relatively low contrast

Partition coefficients capture the compositional contrast across phases. The partition coefficient, *k*, of molecule X is the ratio of its concentrations inside and outside of a condensate, *k* = [*X*]^in^*/*[*X*]^out^ [1, 35, 36]. In live cell experiments, the partition coefficient is estimated as the ratio of fluorescent intensities of a tagged molecule [15, 26, 28–30]. Published values of partition coefficients in live cells range from values close to one to more than a hundred, as shown in Fig. 1a. *In vitro* condensates, on the other hand, tend to have larger values for their partition coefficients, typically on the order of one thousand (Fig. 1a).

**FIG. 1.**
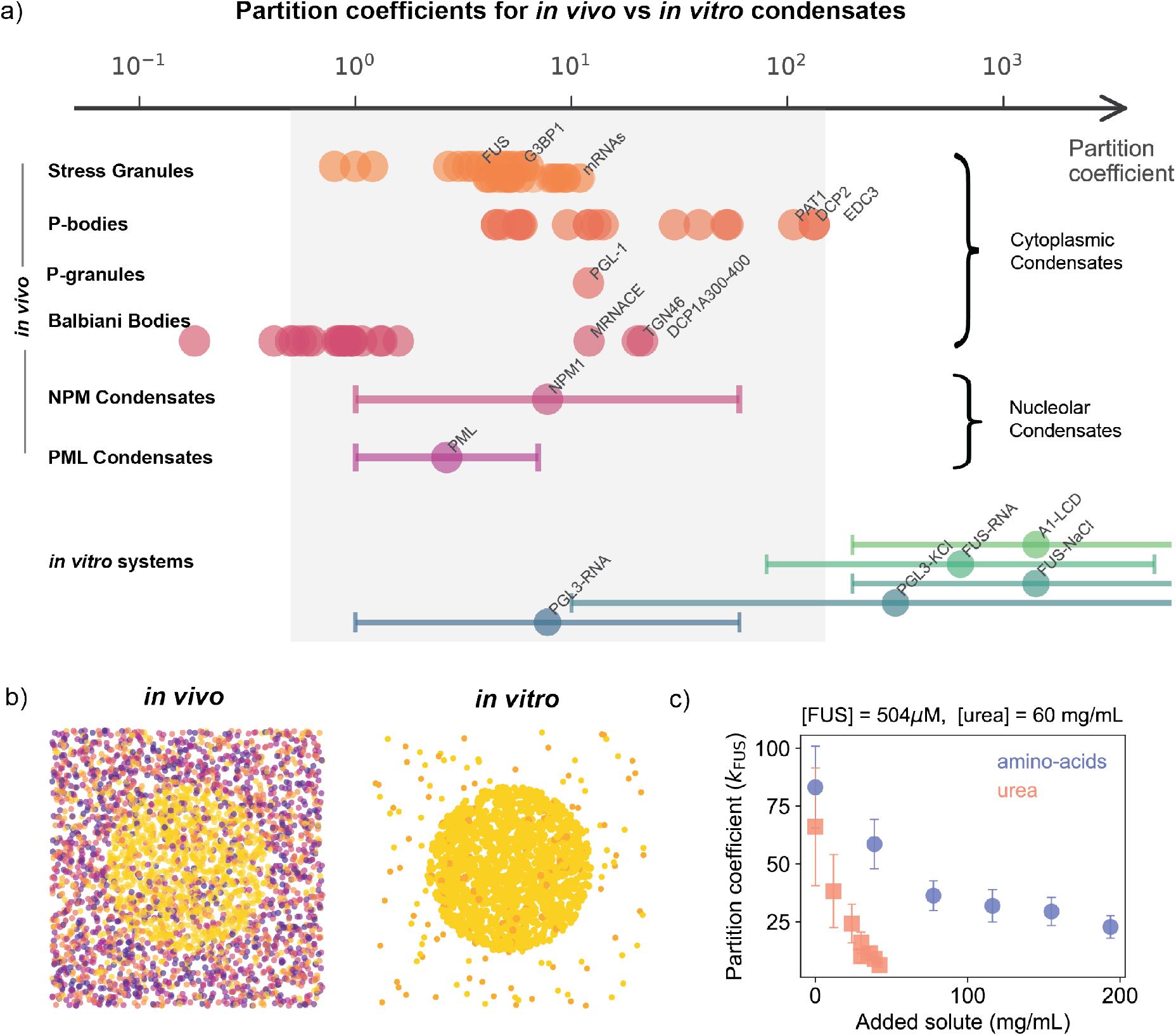
Condensates typically have lower contrasts *in vivo* than *in vitro*: a) Summary of partition coefficients of *in vivo* and *in vitro* condensate systems reported in the literature [15, 26–34] measured using fluorescence techniques. Circles represent measurements of partition coefficients of individual proteins to a particular condensate. Some of the highly partitioning proteins in the condensate have been labeled. The bars on the data indicate the range of partition coefficients that have been measured for the protein due to different expression levels, temperatures or solute compositions. b) Schematic of *in vivo* (left) and *in vitro* (right) condensates with different colors representing different proteins. *in vivo*, some proteins condense, but leave many others in the dilute phase. *in vitro* condensation leaves little protein in the dilute phase. c) FUS partition coefficient, estimated by fluorescence, with addition of a 12 amino acid mixture (blue) or with urea (red) to a solution containing 504*µ*M FUS and 60 mg/mL urea. These fluorescence-based partition coefficients were corroborated by quenching assays and UV-Vis spectroscopy (Supplement S3 D).

This discrepancy between *in vivo* and *in vitro* condensates likely arises from differences in the context of condensation. While *in vitro* systems typically include only drivers of condensation, cellular interiors are filled with thousands of other molecules (Fig. 1b). Generically, this complex crowded environment provides many opportunities for off-target interactions that compete with drivers of phase separation, reducing the effective cohesion between condensate scaffolds [37, 38]. To mimic off-target protein interactions without introducing the complexity of sequence and structure, we titrated a mixture of 12 amino acids into phase-separated solutions of the low-complexity domain of FUS (residues 1-270). As shown by the blue circles in Fig. 1c, the partition coefficient of FUS drops as the samples are supplemented with free amino acids [23]. At a concentration of 200 mg/mL, comparable to estimates of total protein concentration in the cytoplasm [39, 40], we find a four-fold reduction in FUS’s partition coefficient.

Amino acids are not privileged solutes. For example, high concentrations of salt or urea are widely used to solubilize phase-separating proteins during purification [41]. These solutes also have a large effect on the partition coefficient. For example, increasing the concentration of urea from 60 mg/mL to 100 mg/mL reduces the partition coefficient of FUS from from 66 to about 6 ( Fig. 1c).

### B. Low-contrast condensates are more sensitive to solutes

The observed reduction of the partition coefficient with added solute is dominated by changes outside of the condensate. While the concentration of FUS in the dense phase (inside) changes only by a factor of two with added urea, the dilute phase (outside) concentration grows a hundred-fold (Fig. 2a). Considering this large concentration change together with the plentiful and easy-to-purify nature of the dilute phase, it is convenient to quantify the sensitivity of condensates to solutes using the *dilute phase susceptibility, s* ≡ *d*[protein]^dilute^*/d*[solute] [24, 42], *i*.*e*. the slope of the dilute phase protein concentration against the total solute concentration (See inset of Fig. 2b).

**FIG. 2.**
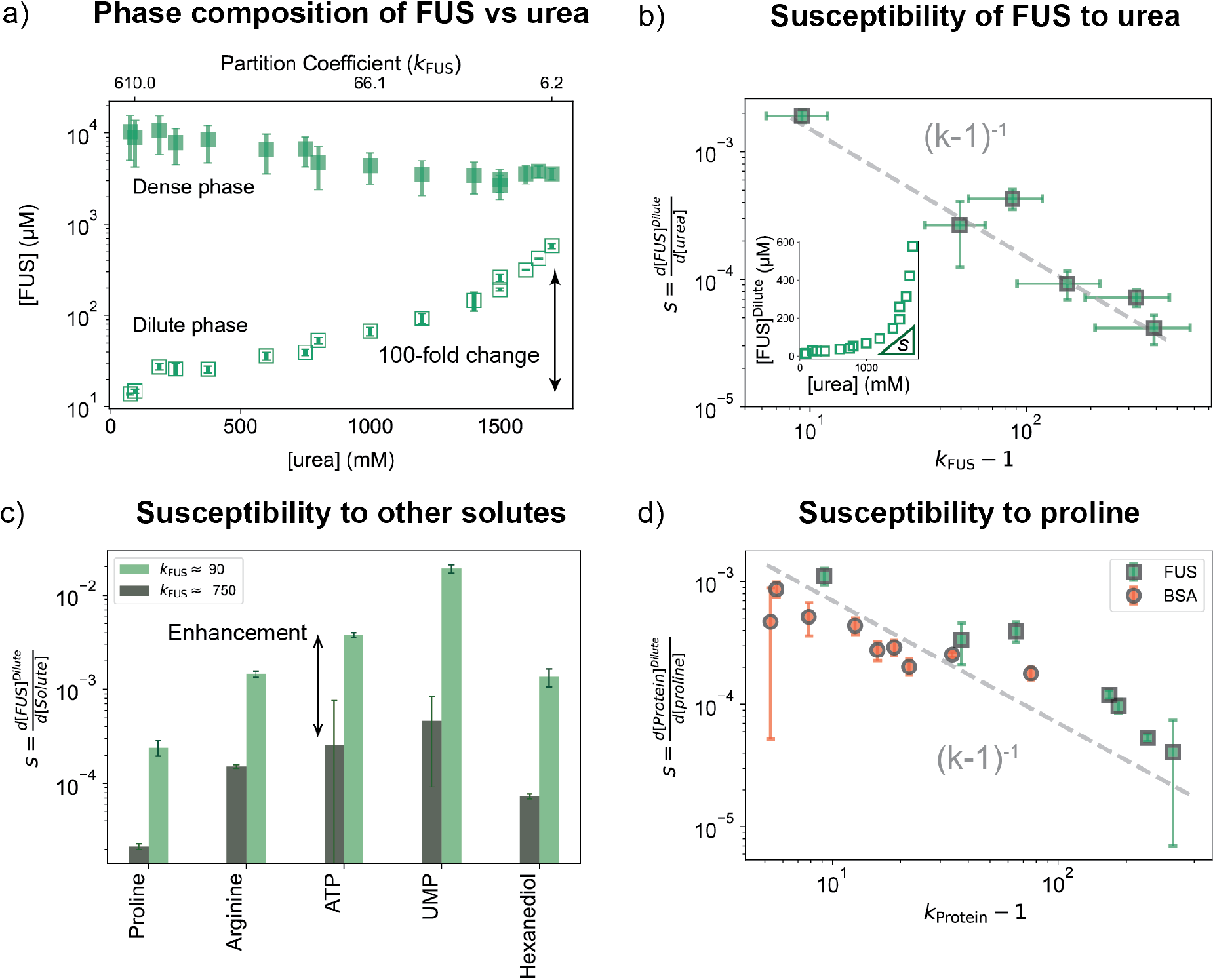
Low-contrast condensates are more sensitive to changes in composition. a) FUS concentration in the dilute (open squares) and dense (filled squares) phases measured with fluorescence as a function of total urea concentration. b) Susceptibility of FUS to urea is defined as the slope of the dilute phase concentration of FUS with respect to total urea concentration (Inset). Susceptibility of FUS to urea increases at lower partition coefficients, roughly following a inverse relationship *s*∼ (*k*_FUS_ −1)^−1^. c) Susceptibility of FUS to other solutes at two different *k*_FUS_ conditions - *k*_FUS_ ≈ 90 (light green) and *k*_FUS_ ≈750 (dark green). d) Susceptibility of FUS and BSA condensates to proline as a function of the protein partition coefficient. (See Supplements S3 C,S2 C for raw data)

FUS condensates are much more sensitive to urea at low contrast. We find that susceptibility increases by a factor of 100 as the partition coefficient (*k*_FUS_) drops from 1000 to 10 (Fig. 2b). Over the range of partition coefficients tested here, the data is consistent with *s*∼ 1*/*(*k*_FUS_ − 1), as shown by the dashed line in Fig. 2b.

FUS’s enhanced sensitivity at low contrast is not limited to urea. We observe this effect across a wide range of molecules, despite differing interactions with the protein. The susceptibility of FUS to two amino acids, two nucleotides, and 1-6 hexanediol are shown in Fig. 2c. At a partition coefficient of 750, the susceptibilities of FUS to these solutes varies from 2 × 10^−5^ to 5 × 10^−4^. After tuning the partition coefficient to 90 using urea, the susceptibility to each of these solutes increases roughly 10-fold. This demonstrates how one solute (in this case, urea) can modulate the response of condensates to other solutes.

The observed enhancement of susceptibility to solutes at low contrast is not exclusive to FUS condensates. For example, condensates of bovine serum albumin (BSA) can be formed by crowding with poly(ethylene glycol) [43, 44]. The partition coefficient of these systems is readily changed by diluting the BSA/PEG mixture with more buffer. We measure the susceptibility of BSA and FUS condensates to proline [23] over a wide range of partition coefficients. While the absolute numbers are different, the susceptibility again varies roughly as 1*/*(*k* −1) in both cases (Fig. 2d), suggesting a general thermodynamic mechanism [24].

### C. Low-contrast condensates are sensitive to thermal fluctuations

So far, we have seen that low-contrast condensates are more sensitive to changes in composition. They are also more sensitive to thermal fluctuations, causing shape [45, 46] and local compositions of the condensate to fluctuate visibly. We first demonstrate this with BSA condensates, tuned from high to low contrast (*k*_BSA_ = 60 to ≈2.5) by dilution with buffer. While the high-*k* droplets were spherical (Supplement S2 E ), the low-*k* droplets display a constantly evolving near-spherical shape, as shown in Fig. 3a, and Supplemental Movie 1. Shape fluctuations suggest ultra-low interfacial tension between the condensate and the surrounding medium [47, 48]. Visible shape fluctuations (*i*.*e*. with an amplitude ≳ 100nm) occur when the interfacial tension is in the order of 0.1 *µ*N/m or lower (Supplement S2 F) [49]. Interfacial tensions of *in vivo* condensates are generally very low, ranging from 0.4 - 70 *µ*N/m, while those of *in vitro* condensates can range anywhere from 0.15 to 4200 *µ*N/m[49, 50].

**FIG. 3.**
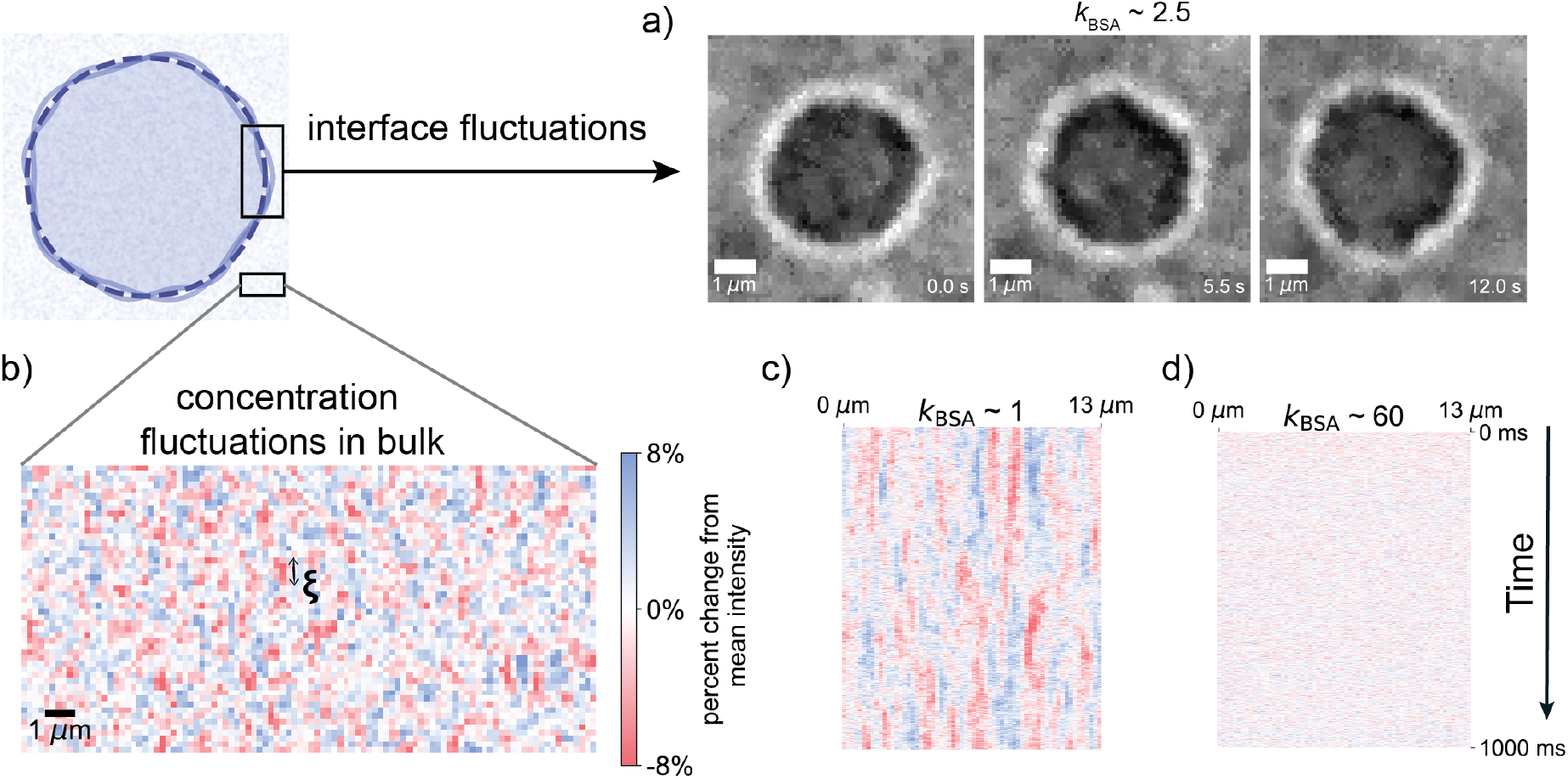
Low-contrast condensates are sensitive to thermal fluctuations. a) Strong shape fluctuations of low-contrast BSA condensates (*k*_BSA_ ≈ 2.5) imaged in bright-field (brightness and contrast adjusted). b) Micron-sized composition fluctuations in the dilute phase *k*_BSA_ ≈ 1 revealed by subtracting mean intensity (averaged over 1 s) from a bright-field micrograph (1 ms exposure). c,d) Persistence of composition fluctuations at *k*_BSA_ ≈ 1 and 60, revealed by bright-field kymographs analyzed as in b. Colorbar is identical in b-d, linear and temporal scales are identical in c and d.

Together with shape fluctuations, both phases of low-contrast condensates also exhibit strong compositional fluctuations [51]. This is suggested by the grainy fluctuating intensity of the bright-field microscopy images in Fig. 3a and Supplemental Movie 1. To make these fluctuations more visible, we acquired a time-series of bright-field microscopy images, and subtracted off the time-averaged intensity at each pixel and timepoint. For the dilute phase of BSA/PEG and *k* ≳1, we observe fluctuations in light intensity (Fig. 3b and Supplemental Movie 2). Intensity fluctuations have a standard deviation of about 2.5%, a characteristic length scale of about one micron, and persist for about 200 ms, as shown by the kymograph in Fig. 3c. Large intensity fluctuations imply that there is a small energetic penalty to concentrate the protein, as captured by the osmotic compression modulus, *K* [52, 53], here estimated to be of order kPa (see Supplement S2 G). When the contrast between the phases is larger, these fluctuations are no longer resolvable, as shown by Fig. 3d.

We observe similar results for condensates formed by Bik1, a yeast protein associated with the plus-end of microtubules [54]. In this system, we achieve low partition coefficients by increasing the NaCl concentration. At low contrast (*k*_Bik1_ *<* 2), Bik1 condensates show shape and composition fluctuations similar to those of BSA condensates (Supplement S4 B and Supplemental Movie 3). We cannot reliably measure partition coefficients of Bik1 below two because laser exposure shifts the phase equilibria, as demonstrated in Supplemental Movie 4. This response further emphasizes the sensitivity of low-contrast condensates to external perturbations.

### D. Low-contrast condensates have distinct structure and properties

Shape and composition fluctuations seen in Fig. 3 suggest that interfacial tensions and osmotic compression moduli vanish at low contrast.

To quantify interfacial tension, *γ*, we bring together two dense-phase droplets with holographic optical tweezers and image their fusion with a high speed camera (inset Fig. 4a) [19, 55–57]. We locate droplet edges, and quantify their relaxation from ellipsoidal to spherical shapes (see Supplement S2 H). The time to fuse, *τ*, is proportional to the final droplet radius *τ* = *R*_final_*/v*, where *v* is the capillary velocity [58, 59], which depends on the interfacial tension and viscosities of the two phases. The capillary velocity varies non-monotonically with *k*_BSA_, with a peak value of 110 *µ*m/s at *k*_BSA_ ≈ 10, as shown by orange circles in Fig. 4a.

**FIG. 4.**
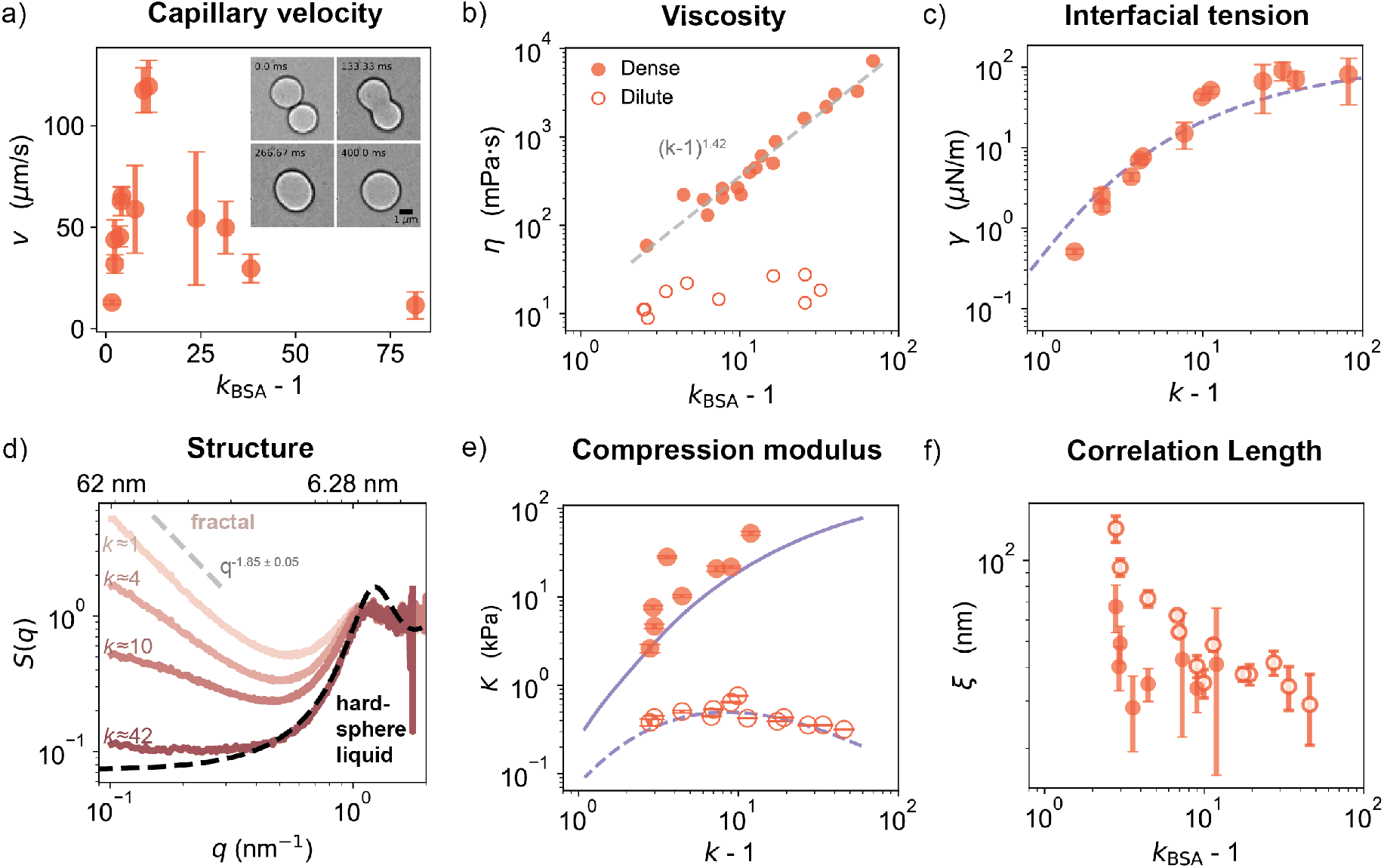
Variation of condensate structure and properties with partition coefficient. a) Analyzing droplet merger events (inset), capillary velocity is measured for droplets with varying partition coefficients of BSA (*k*_BSA_). For each partition coefficient, 16 to 24 droplet mergers are analyzed. b) Viscosity of the dense phase (filled circles) and dilute phase (open circles) as a function of partition coefficient. The dashed gray line shows a fit to the dense phase data using a power law. c) Interfacial tension as function of partition coefficient of BSA combining viscosity and capillary velocity data. The blue dashed line is the scaled interfacial tension of the liquid-vapor interface of water plotted against the partition coefficient of water. d) SAXS structure factor of dense phase of BSA condensates for four different values of *k*_BSA_. The dashed black line is the theoretical Percus-Yevvick structure factor for hard sphere liquids with volume fraction 0.33. For *k ≈*1, the long wavelength behavior was fit to a power law and the exponent is reported. The dashed gray line is a guide to the eye. e) The osmotic compression modulus of the dilute (open circles) and dense phase (filled circles) obtained from the structure factor as a function of the BSA partition coefficient. The scaled isothermal compression modulus of liquid (solid blue) and vapor (dashed blue) phases of water are shown against the partition coefficient of water. f) The characteristic size of concentration fluctuations (correlation length) obtained from the structure factor for the dilute (open circles) and dense phases (filled circles).

To extract interfacial tension from capillary velocity, we need to quantify the viscosities of the dense and dilute phases. In the limit of high viscosity-contrast, *v* = *γ/η* [58], where *η* is the viscosity of the dense phase. For a more general formula, including low viscosity contrast see Supplement S2 H 3. We measure the viscosity of the dense and dilute phases using a rheometer (see Fig. 4b and Supplement S2 I). The viscosity of the dense phase (filled circles) increases one hundred-fold from 50 to nearly 10,000 mPa · s while that of the dilute phase (open circles) stays relatively constant. This dramatic increase in viscosity at large *k*_BSA_ occurs as BSA approaches close packing in the dense phase (see Supplement S2 K) [60, 61] Interfacial tensions of BSA condensates obtained from the viscosity and capillary velocity data are shown in Fig. 4c. At high contrast (*k*_BSA_ *>* 20), the interfacial tension is around 100 *µ*N*/*m and is largely insensitive to partition coefficient. As the contrast is lowered, the interfacial tension drops by more than two orders of magnitude, reaching 0.3 *µ*N*/*m at *k*_BSA_ = 2.5.

This drop of interfacial tension at low contrast is generic for two-phase systems. Interfacial tension quantifies the energy penalty of two phases coming into contact. Intuitively, this must vanish as the two phases become identical (*k* →1). To highlight the ubiquity of this effect, we superimpose scaled values of the liquid-vapor interfacial tension of water (dashed blue line) upon the BSA-condensate data in Fig. 4c [62, 63]. The functional forms of the decay in these very different systems are remarkably similar: each vanishes as *k* → 1 and plateaus beyond *k* ≈ 10.

To quantify the osmotic compression modulus, *K*, we use Small Angle X-ray Scattering (SAXS). SAXS provides the structure factor, *S*(*q*), which quantifies the magnitude of concentration fluctuations as a function of wavelength, 2*π/q* (Supplement S2 J) [64, 65]. In the limit of long wavelengths (low *q*), the osmotic compression modulus is related to the structure factor: *K* = *ρk*_*B*_*T/S*(*q*→ 0), where *ρ* is the number density [52, 64]. Structure factors of dense-phase BSA condensates are shown for a range of partition coefficients in Fig. 4d. As the partition coefficient drops, *S*(*q*) increases at low *q*. To extract the compression modulus from this, we fit the low *q* behavior (*q <* 0.25 nm^−1^) to the following scattering form [66] (Supplement S2 J):

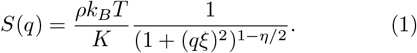

The osmotic compression modulus of the dense phase (closed circles in Fig. 4e) drops from 50 to 1 kPa as *k*_BSA_ decreases from 10 to 2.5. The dilute phase modulus (open circles) is lower, around 0.3 kPa, and is relatively independent of partition coefficient. As for the interfacial tension, these trends are generic for a low-contrast two-phase system. For further comparison, we superimpose scaled isothermal compression moduli of coexisting liquid (solid blue line) and vapor (dashed blue line) phases of water upon the BSA-condensate data in Fig. 4e (See Supplement S5 B and [62, 63]) The moduli of the dense and dilute phases of these very different systems show remarkably similar variation with partition coefficient.

The shape of the structure factor conveys the organization of molecules in solution. A perfectly random organization (equivalent to an ideal solution) would result in a uniform structure factor, *S*(*q*) = 1. Liquid-like structures feature a drop at low-*q*, and peaks reflecting the intermolecular spacing. At high contrast (*k*_BSA_ ≈ 40), the structure factor of the dense phase is liquid-like, roughly following the predicted scattering [67] from a hard sphere liquid with volume fraction 0.33 (dashed black line). At lower contrasts, the structure factor curves up at low-*q*, approaching a power law (straight line on a log-log plot). This indicates that the BSA molecules are arranged into a fractal (*i*.*e*. scale-invariant) structure [68, 69]. At the lowest contrast, the fractal structure spans the widest range of wavelengths. The fractal dimension at low *q* is found to be 1.85 ± 0.05 *i*.*e. S*(*q*) ∼*q*^−1.85*±*0.05^. For comparison, this value is between the fractal dimensions of polymers in a theta and good solvents (2.0 and 1.7, respectively) [70]. The scattering at larger *q* values (*q* ≳ 1 nm^−1^) does not significantly change with partition coefficient, implying that the protein conformation is relatively stable.

The characteristic size of compositional heterogeneities is revealed by the correlation length, *ξ*, from the fit of *S*(*q*) to Eq. 1. At the lowest partition coefficients, these fluctuations are large enough to be seen under the microscope, as shown earlier in Fig. 3b. However, SAXS resolves concentration fluctuations at much smaller-wavelengths, as shown in Fig. 4f. The correlation length in both the dilute (open circles) and dense (closed circles) phases are comparable and have similar trends. The correlation length in the dilute phase increases from *ξ ≈* 10 nm at *k*_BSA_ ≈10 to more than 100 nm at *k*_BSA_ ≈ 3. The latter is more than 10 times the nearest neighbor spacing between protein molecules at that concentration (∼8 nm).

### E. A unified description of condensates based on partition coefficient

Large correlations and fractal structure also appear in other two-phase systems when the contrast between the phases is low [71]. When correlations are large compared to the size of the molecules, they dictate material properties. Very different systems, therefore, can behave similarly despite molecular differences. This is known as *universality* and is well-established in the behavior of magnets, fluids, and other systems with two coexisting phases [71–76].

Across these diverse systems, universal behavior is dictated by proximity to a *critical point* where the contrast between the two phases completely vanishes. Properties studied in the previous section have well-known scaling relations with the ‘distance’ to the critical point, *t*, also known as the normalized temperature. Close to the critical point, the interfacial tension, *γ*, isothermal compression modulus, *K*, and correlation length, *ξ*, scale with *t* as *γ* ∼ *t*^*µ*^, *K* ∼ *t*^−*γ*^, and *ξ* ∼ *t*^−ν^, respectively. The exponents are predicted to be *µ* = 1.29, *γ*^, *≈*^ 1.31 and ν = 0.629. [66, 74, 77, 78] To apply these predictions to multi-component condensates, we need to determine an analogous normalized temperature for our isothermal systems. To do so, we turn to phase diagrams.

Consider the liquid-vapor phase equilibrium of water shown in Fig. 5a for which these ideas are well established [63]. The vapor phase (open triangles) meets the liquid phase (closed triangles) at a unique temperature (*T*_*C*_) and water concentration - the *critical point* (red star). The gray lines are called ‘tie lines’ and connect every vapor composition to its coexisting liquid phase, here at the same temperature. Quantifying the distance to the critical point with the normalized temperature *t* = |(*T* −*T*_*C*_)*/T*_*C*_|, the shape of the phase diagram can be directly compared to the predictions of the 3D Ising model, originally developed for ferromagnets [79, 80]. This prediction (described in the Supplement S5 A) agrees well with data near the critical point, as shown by the dashed black line in Fig. 5a. The 3D Ising model also accurately captures the temperature dependence of water’s interfacial tension and isothermal compression moduli (Supplement S5 C).

**FIG. 5.**
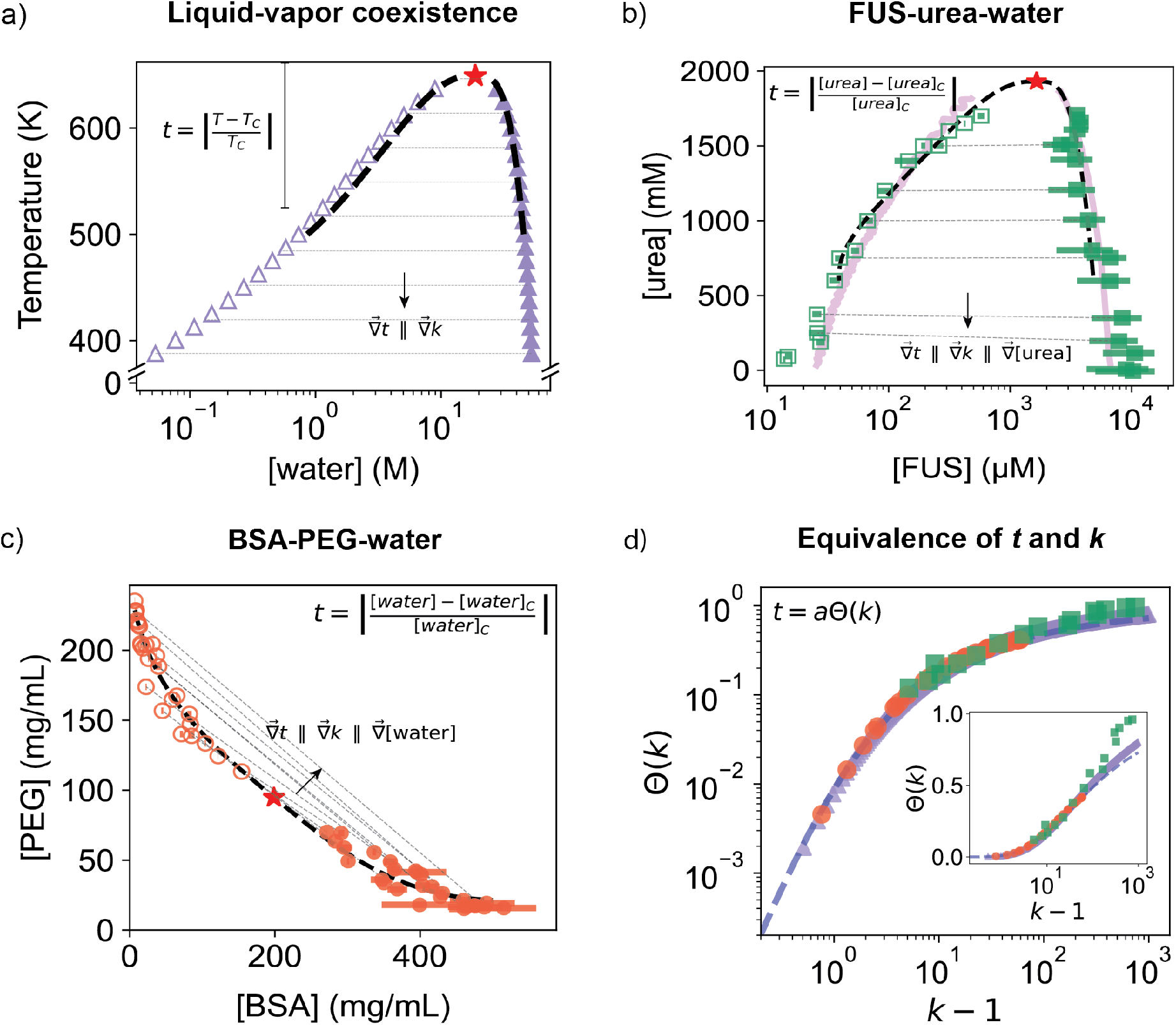
Connecting temperature- and concentration-driven phase behavior. a) Liquid (filled blue triangles) - Vapor (open blue triangles) coexistence of water from NIST database [62, 63] and the fit to 3D Ising model (thick dotted black line) for low partition coefficients (*k*_water_ *<* 100) using the normalized temperature variable *t* shown in the figure (Supplement S5). The red star shows the critical point and the thin dotted gray lines are tie lines. The gradient of the partition coefficient of water is perpendicular to the tie line and away from the critical point. b) Coexistence of dense (filled green squares) and dilute (open green squares) phase of the ternary system of FUS, urea and water. The thick dotted black line is the 3D Ising model fitting form. The red star is the critical point predicted from fit. The pink curve is the phase boundary given by Flory Huggins theory (Supplement S3 F). c) Coexistence of dense (filled orange circles) and dilute (open orange circles) phase of the ternary system of BSA, PEG and water. Again, the thick dotted black line is the fit of the data with a 3D Ising model generalized to ternary systems (Supplement S2 L). d) All three model systems show the same one-to-one relationship between the partition coefficient and normalized temperature. Dashed line shows the empirical fit to Θ(*k*) = (1 − *k*^−1*/*3^)^3^.

To apply critical point concepts to isothermal multi-component systems, we quantify distance from the critical point in terms of concentrations [75]. In these cases, tie lines still connect coexisting phases, but can have arbitrary orientations, as shown by the the isothermal phase diagrams of FUS-urea and BSA/PEG in Fig. 5b-c. Along each tie line, the partition coefficient of every component is unchanged. Moving perpendicular to the tie lines brings you toward or away from the critical point.

In the case of FUS-urea, tie lines are oriented in the direction of constant urea concentration (Fig. 5b) as verified by Raman spectroscopy (Supplement S3 E). Assuming urea concentration plays the role of temperature, we fit the FUS-urea data to the 3D Ising model using *t* = |([urea] − [urea]_C_)*/*[urea]_C_ |, where [urea]_C_ is the concentration of urea at the critical point. As shown by the dashed black line in Fig. 5a, we find reasonable agreement for *k*_FUS_ *<* 100, comparable to predictions of molecularly motivated multi-component Flory-Huggins theory (pink curve) with fixed *χ* parameters [81–83]. See Supplement S3 F and S3 G for details.

In the case of BSA/PEG, tie-lines point along neither axis (Fig. 5c). We fit for the phase boundary using the orientation and magnitude of *t* as free parameters. The resulting fit, shown as a black dashed line, nicely matches the data over the entire experimental range and identifies the *t* direction as roughly perpendicular to the measured tie lines. Accounting for the molar volumes of PEG and BSA, we find that fitted *t*-direction is well-aligned with the direction of increasing water concentration: *t ≈* |([water] [water]_C_)*/*[water]_C_ |, with [water]_C_ being the concentration of water at the critical point (Supplement S2 L). The scaling of the interfacial tension, isothermal osmotic compression moduli, and correlation length with *t* for this system appear to be consistent with predicted scalings (Supplement S2 M).

The above results suggest that the theory of critical phenomena could provide a unified framework for fitting phase diagrams and predicting material properties. As typically formulated, implementing this approach would require that one measures and fits a phase diagram (with tie lines) to determine the critical composition and normalized temperature direction. Alternatively, our data suggest that partition coefficients of condensate scaffolds can serve as a proxy for the normalized temperature. Specifically, Fig. 2b-d found that the solute susceptibilities scaled as *s* ∼1*/*(*k* −1) for all tested solutes and scaffolds. Fig. 4c-e showed remarkable similarities in the scaling of material properties of BSA/PEG system to the liquid-vapor coexistence of water. To refine this hypothesis, we plot fitted values of *t* and measured values of *k* across the phase diagrams of water, FUS, and BSA (Fig. 5d). Strikingly, all three systems collapse to a single curve, Θ(*k*), when normalized by a system dependent scale factor, *a*. Empirically, Θ(*k*) ≈ (1 −*k*^−1*/*3^)^3^ (dashed line in Fig. 5d). This allows one to easily predict how material properties change with *k*. Given two condensates of the same type with different *k* values, this approach can readily predict relative values of interfacial tension and osmotic compression moduli (*e*.*g. γ*_2_*/γ*_1_ = (Θ(*k*_2_)*/*Θ(*k*_1_))^*µ*^ ). Importantly, this approach could be readily extended to condensates in live cells.

## CONCLUSIONS

Biomolecular condensates tend to show lower protein enrichment (or partition coefficient) in cells than in typical *in vitro* models. We hypothesize that low contrast is a natural consequence of off-target interactions that compete with drivers of condensation. Consistent with this, a mixture of free amino acids reduces the partition coefficient of FUS by a factor of four. This observation echos recent results on the impact of metabolites and small molecules on the phase equilibria of biomolecular condensates [18, 23, 24, 36, 84]. As contrast vanishes, condensates become more sensitive to their surroundings and develop strong shape and concentration fluctuations. In this limit, systems with very different molecular structures and interactions (*e*.*g*. liquid-vapor coexistence of 18 Da water and liquid-liquid phase separation of 66 kDa BSA) have similar trends in their material properties. These results can be quantitatively understood by expressing the theoretical framework of critical phenomena [45, 79, 80] in terms of partition coefficients.

These results have broad implications for the study of critical fluids. Typically, distance from the critical point is measured using the normalized temperature, *t*. Our results suggest that *k*− 1 is superior from an experimental point of view, as distance from the critical point can be assessed from a single concentration ratio with no other knowledge of the phase diagram. This experimental convenience does not sacrifice predictive power: *k* −1 is at least as effective as *t* at capturing the universal scaling of material properties across diverse two-phase fluid systems (See Supplement S6 B). On the other hand, further theoretical work is required to generalize *k* −1 to systems with many components, and account for multi-phase coexistence, diverse universality classes [85], and activity [86–89].

The generic behavior of condensates at low contrast has implications for understanding the material properties of condensates within cells. Since condensate material properties can change by orders of magnitude as the contrast between phases disappears (Fig. 4), *in vitro* experiments quantifying condensate material properties [11, 19, 22, 50, 90] should take care to match partition coefficients with cellular values. Further, live cell measurements of partition coefficients [15, 26, 28–30] could reveal dynamical changes to material properties through the universal scaling of material properties described in Section I E.

The unusual properties of low-contrast condensates could enable cellular physiology. Stress granules’ low contrast (see Fig. 1a) amplifies their sensitivity to biological stress (see Fig. 2c) [91]. The low interfacial tension of low-contrast condensates facilitates their spreading on cytoskeletal filaments and cellular membranes [5, 92–95]. Low osmotic compression moduli (S 10kPa) suggest new mechanisms for mechanobiological feedback, where condensate phase equilibria are perturbed by cytoskeletal forces [7]. Finally, material properties of low-contrast condensates can be readily tuned over a wide range through biochemical activity, enabling post-translational regulation of condensate function.

## II. ACKNOWLEDGEMENTS

We thank William Jacobs, Ned Wingreen, Andrew Pyo, Kathryn Rosowski, and members of the Dufresne lab for helpful discussions and comments on the manuscript. We also thank Anna Barth and Itai Cohen for helpful discussions and access to their rheometer. This work was partially supported by the Swiss National Science Foundation through NCCR Bio-Inspired Materials (Grant No. 205603) and the Sinergia program (Grant No. 189940). TM was supported by a Schmidt Science Fellowship in partnership with Rhodes Trust and CL was supported by the Deutsche Forschungsgemeinschaft (Proj.-670Nr. 523842861, LO 3088/1-1). This work made use of the Cornell Center for Materials Research shared instrumentation facility. This work is based on research conducted at the Center for High-Energy X-ray Sciences (CHEXS), which is supported by the National Science Foundation (BIO, ENG and MPS Directorates) under award DMR-2342336, and the Macromolecular Diffraction at CHESS (MacCHESS) facility, which is supported by award 1-P30-GM124166 from the National Institute of General Medical Sciences and the National Institutes of Health. This work made use of the Cornell University NMR Facility, which is supported, in part, by the NSF through MRI award CHE-1531632. This work benefited from measurements on a BioXolver X-ray Source from Xenocs, supported by NIH grant S10 OD028617.

## III. AUTHOR CONTRIBUTIONS

K.V. and E.R.D. designed the project, analyzed, and interpreted the results, and wrote the manuscript with inputs from T.M, S.B-D, C.L, T.B, R.W.S, S.J.T, S.D. and J.P.S. K.V. performed all of the experiments except for dense phase rheology (B.S. and T.M.) and some Raman and susceptibility data (D.M.) K.V and T.B purified proteins with assistance from C.L.

## SUPPLEMENTAL INFORMATION

### S1. MATERIALS

#### A. Purchased Materials

Bovine serum Albumin (BSA) (A7638), L-Isoleucine (I2752), L-Aspartic acid (A9256), L-Alanine (A7469), L-Threonine (T8625), D-Glutamine (G9003), L-Serine (S4500), Glycine (G7126), L-Leucine (L8000), L-Glutamine (G5792), Urea (U5128), Potassium phosphate dibasic trihydrate (P5504), HEPES (H23830), Helmanex III (Z805939), tetramethylethylenediamine (TEMED, T9281), acrylamide (A8887), PEGMA (poly(ethylene glycol) methyl ether acrylate, 454990), trimethoxysilyl-propylmethacrylate (TOPA, M6514), ammonium persulfate (APS, A3678) were purchased from Sigma-Aldrich.

Polyethylene glycol 4000 (A16151), Alexa Fluor 488 NHS Ester, succinimidyl ester (A20000) was purchased from Thermo scientific. Potassium Chloride (P217), Potassium Phosphate Monobasic (P285), Potassium Chloride (S271) and 1,6-Hexanediol (A12439.30) was purchased from Fisher chemical. L-Arginine (ALX-101-004-G025) was purchased from Enzo Life Sciences. DL-Glutamic Acid (Cas 19285-83-7) Lysine Hydrochloride (L1142), L-Proline (PR120) was purchased from Spectrum Chemical MFG Corp. Deuterium oxide (DLM-4) was purchased from Cambridge Isotope Laboratories. All water used is milli-q.

#### B. FUS Purification

His_10_–Gb1–TEV–FUS (residues 1–270) was overexpressed in *E. coli* BL21 (DE3) (C2527H, New England Biolabs) with 0.2mM IPTG and 20 ^°^C overnight. Cells were lysed with an ultrasonic homogenizer and centrifuged at 5000g for 25 mins at 4^°^C to isolate the inclusion bodies. The pellet is resuspended in buffer (8 M urea, 50 mM HEPES, 500 mM NaCl, pH 7.5) and dounced to disrupt the inclusion bodies. The lysate is then clarified by centrifugation at 16000g for 25 mins at 4^°^C. The protein was isolated using nickel affinity chromatography with an elution buffer (1 M urea, 50 mM HEPES, 150 mM NaCl, 500mM Imidazole, pH 7.5). The Gb1 tag is cleaved using His_10_–TEV protease (in-house, pET29b-10xHis-Super TEV [96]) and the solution was dialyzed overnight to 1 M urea, 50 mM HEPES, 150 mM NaCl and 1 mM DTT, pH 7.5. The protein is then dialyzed into the storage buffer (6 M urea, 50 mM HEPES, 150 mM NaCl, pH 7.5) which prevents phase separation when the protein is concentrated. The solution is put through the nickle affinity column again to remove all the cleaved His tags. The purified protein is then concentrated to around 82 mg/mL, snap frozen and stored at -80^°^C.

#### C. Bik1 Purification

N-terminal His_6_–HRV3C–Bik1 was over-expressed in *E. coli* BL21-CodonPlus (DE3)-LOBSTR (230280, Agilent) with 0.75mM IPTG at 20 ^°^C overnight. Cells were lysed using ultrasonic homogenizer and centrifuged at 16000g for 25 mins at 4 ^°^C. The protein was purified by nickel affinity and size-exclusion chromatography in buffer containing 50 mM Tris, 500 mM NaCl, 1 mM DTT, and 5% (v/v) glycerol, pH 7.5. Once purified, the protein was concentrated to a concentration of 731.6 *µ*M, snap frozen and stored at -80^°^C.

#### D. PEGMA–acrylamide glass coating

All glass slides and coverslips used for imaging in the manuscript were coated with PEGMA-acrylamide glass coating to prevent the wetting of condensates to the glass. Glass slides and coverslips were first cleaned with 1% Helmanex III and boiling water, sonicated for 3 mins and rinsed. They were then sequentially treated with 200 proof ethanol and 100 mM KOH, with 3 minutes of sonication in each solution, followed by rinsing after each step. They were silanized with trimethoxysilyl-propylmethacrylate (TOPA). They are then coated with 1.5% PEGMA (poly(ethylene glycol) methyl ether acrylate) and 0.5% acrylamide solution polymerized with ammonium persulfate (APS) and tetramethylethylenediamine (TEMED) and stored in air tight containers.

### S2. BSA/PEG SYSTEM

#### A. Condensate Preparation

For the BSA/PEG condensate system, stock solutions of each individual constituent is prepared at a high concentration in water: BSA at 360 g/L, PEG(4k) at 400 g/L, Potassium Chloride (KCl) at 3.5M and potassium phosphate buffer (KP) at 1M. The concentration of BSA in the stock solution is measured using the Nanodrop One UV-Vis spectrometer after it has been prepared. BSA and PEG(4k) stock solutions are stored at 4C. For making BSA/PEG condensates, appropriate volumes of water, KCl, KP buffer and BSA are mixed first. Then, the appropriate volume of PEG(4k) is added and mixed thoroughly by vortexing, triggering the formation of condensates. Dilute and dense phases of phase-separated solutions of BSA and PEG are isolated by centrifuging the sample at 16900g and 23^°^C using the Eppendorf 5418R centrifuge. Centrifuge times varied from 10 mins to 1 hour depending on the density difference between the two phases.

#### B. Measurement of BSA concentration

The concentration of BSA in the dense and dilute phases of BSA/PEG condensates is measured using the absorbance of the solution at 280nm with the Nanodrop One UV-Vis spectrometer. The solution is appropriately diluted with water so that the absorbance at 280nm lie in the range from 0.05-10 A.U. The extinction coefficient for BSA used is 43.8M^−1^cm^−1^.

#### C. Susceptibility measurements

For the preparation of BSA/PEG condensates containing proline, appropriate volumes of water, KCl, KP buffer, proline and BSA are thoroughly mixed. Then, the appropriate volume of PEG(4k) is added and mixed thoroughly by vortexing, triggering the formation of condensates. Five such phase-separating mixtures whose average composition varies only in the proline concentration are prepared. Dilute phase is isolated by centrifuging the sample at 16900g and 23^°^C between 10 mins to 30 mins. BSA concentration in the dilute phase is measured as described in the previous section. A straight line is fit through the dilute phase BSA concentration vs proline concentration and the slope is reported as susceptibility. Raw curves of the susceptibility of BSA to proline are shown in Fig. S1.

**FIG. S1.**
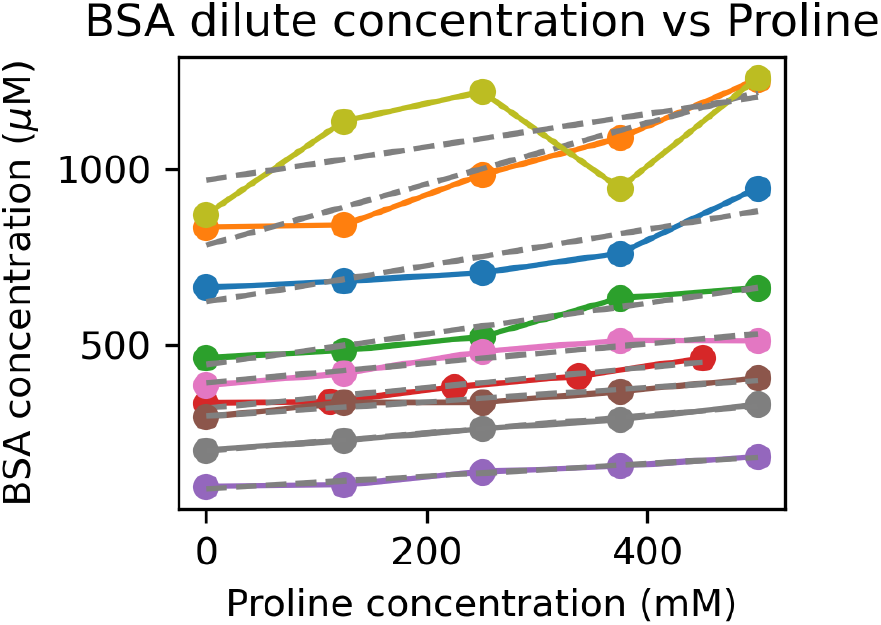
Susceptibility of BSA to proline

#### D. Measurement of PEG concentration

As done in [43], we use NMR spectroscopy to measure PEG(4k) concentration. To an appropriate volume of sample (50*µ*L for dense phase or 100*µ*L for dilute phase), we add enough D_2_O to increase the volume up to 550*µ*L. Then we add 50*µ*L of DMF as a reference molecule and thoroughly mix the sample by vortexing. We obtain the NMR signal using the Av400 Bruker NMR instrument, averaging over 4 scans with a 30s relaxation decay. The data was phase and background corrected using Mestrenova software. Integrals were evaluated using python. We can identify the DMF peak in the NMR spectrum and since it is of known concentration, compare the area of the peaks of DMF to the peaks associated with PEG(4k) in order to compute the molar concentration of PEG(4k).

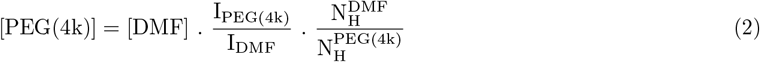

where, [X] is the molar concentration of the molecule, I_X_ is the area under the peak for X and 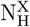 is the number of hydrogen atoms associated with that peak in each molecule (X = DMF,PEG(4k)).

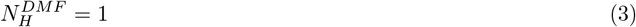

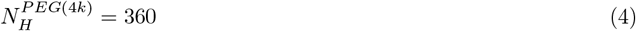

In order to test the validity of this method, we took PEG stock and diluted it until concentration of 90 g/L - which is close to the PEG concentration at the critical point. Then we made 5 independent measurements of this sample using the above method and measured the PEG concentration with 0.5% error as shown in Fig. S2 c). We also made 3 independent measurements of the dilute and dense phase for *k*_BSA_ = 10. We estimate about 2.5% error in each of these samples.

**FIG. S2.**
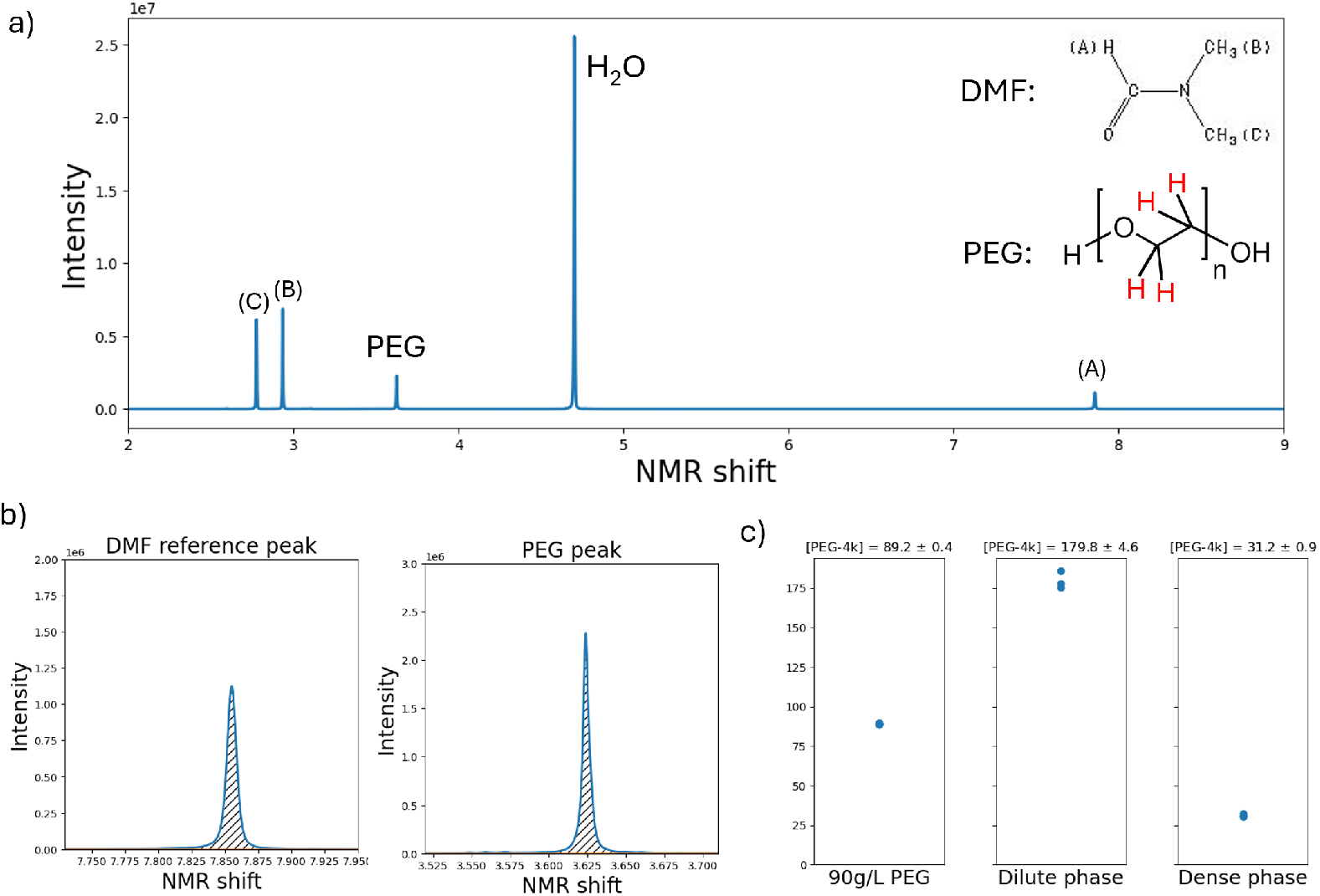
NMR measurements and error analysis for PEG concentration. a) NMR spectrum of solution containing dilute phase and DMF reference. The peaks corresponding to each H atom in the molecule is shown. b) DMF and PEG peaks used for measuring concentration of PEG. The area under the peak (shown in stripes) is integrated. c) Estimating error bars of measurement of PEG using NMR. The dilute and dense phase measurements show an error of about 2.5%.

#### E. Shape of BSA/PEG droplets at high partition coefficients

As shown in supplementary Fig. S3, BSA/PEG droplets at large partition coefficients (*k ≈*60) have robust spherical shapes at all times.

**FIG. S3.**
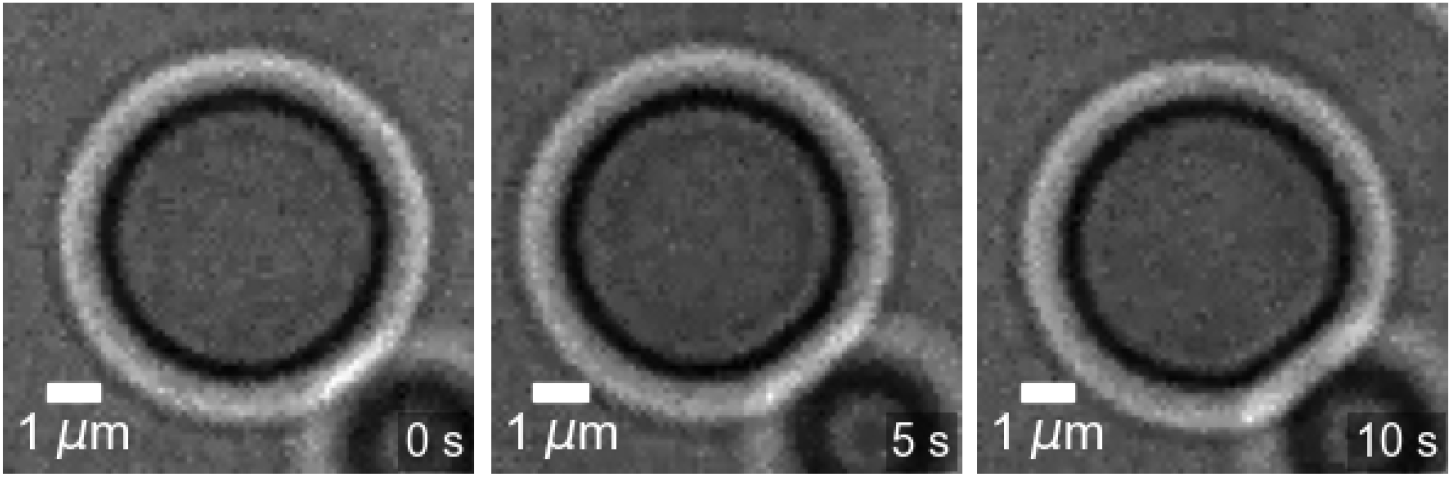
BSA/PEG droplets have robust spherical shapes at high partition coefficients (brightness and contrast adjusted)

#### E. Estimating interfacial tension for visible fluctuations

We observe visible fluctuations of the interface when the energy to excite capillary waves on the surface of the condensates is comparable to *k*_*B*_*T* . The waves are decomposed into spherical harmonics *Y*_*lm*_ and the energy of each wave is calculated [97]. Using the equipartition theorem, the thermally averaged amplitude of each mode is computed. This is then summed to produce the mean square amplitude of fluctuations from mean radius of the droplet.

This is found to be:

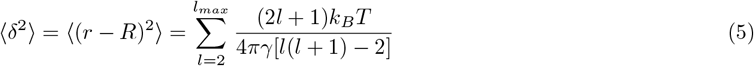

where *R* is the average radius of the droplet and *l*_*max*_ is the largest mode that can be resolved with the instrument. In order to observe a change of 100nm (δ = 100nm) on the surface of a droplet, we choose *l*_*max*_ = 2 giving *γ* ≈ 0.3 × 10^−7^N/m.

#### G. Estimating the osmotic compression modulus from intensity fluctuations

The osmotic compression modulus is estimated using the formula:

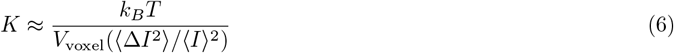

where *V*_voxel_ is the volume of each voxel.

We observe intensity fluctuations of ≈2.5%. The image was taken with a 100X Oil Immersion objective who’s point spread is ≈200nm - This covers about 2 pixels. The standard deviation of 2×2 binned intensities is 2%. We estimate the depth of the field of view to be in the order of the wavelength of light 500nm. This gives us an estimate of the volume of the voxel as *V*_voxel_ ≈ 200nm × 200nm × 500nm ≈ 2 × 10^−20^m^3^.

Plugging this into our expression of *K*, we obtain *K* ≈ 0.5kPa.

#### H. Capillary velocity measurement

Fig. S4a) provides a schematic of the holographic optical trap setup. The laser for the optical trap is the Omicron LuxX 785-200 (200mW, 785nm wavelength). Using an SLM (Meadowlark Optics, S19X12-500-1200-HDM10), we can control the position and number of optical traps in the field of view using python code in real time. A 60x water-immersion objective (NA 1.3) was used for all measurements. Images of droplet fusion are taken with the FastCam Mini AX200. The frame rate is chosen appropriately (between 250 - 1000 fps) such that there is at least 200 frames during the fusion event.

Pairs of droplets are lifted off the coverslip by moving the focal plane of the objective above and away from the coverslip. This ensures that interfacial effects from the wetting of droplets on to the coverslip do not affect the measurement. Care is taken to bring both droplets to the focal plane of the microscope by individually adjusting individual optical traps in the z-direction. The droplets are then slowly moved closer to each other in logarithmically spaced steps. Once they are close enough, the traps are stopped and droplets eventually merge when they touch each other through Brownian motion.

Fig. S4b) shows the fusion event of a representative droplet pair. Ignoring the initial necking behavior, we only analyze the shape of the droplet at late times as it relaxes from an ellipsoidal to spherical shape. For each sample 16-24 droplet merger events are recorded and analyzed.

##### 1. Shape analysis of the droplet

To find the boundary of the droplet, we first find an approximate center of the droplet using its intensity weighted “center of mass”. Then a radial gradient is computed from this approximate center. There will be two large gradients at the interface of the droplet - a large negative gradient from the inside of the droplet to the dark edge and a large positive gradient from the dark edge to the outside of the droplet. We perform a multi Otsu thresholding [98] with three classes to capture the large negative, large positive and the zero gradient regions. We choose specifically the large negative class and this selects the inner ring of the droplet. Once we obtain all the pixels identified as the inner boundary of the droplet, we fit it to an ellipse to obtain the major (*L*) and minor axes (*W* ) lengths of the ellipse (shown in S4c)

##### 2. Fitting the shape parameter as a function of time

From the major and minor axes of the ellipse, a shape parameter, *A*, is computed. This is then fit to an exponential decay with a constant positive offset.

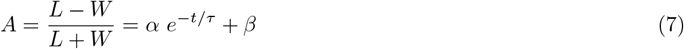

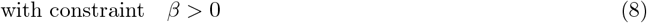

We are mainly interested in the merger timescale *τ* . Fig. S4d) shows the fit of the shape parameter for one droplet taken at a given composition (*k* = 5.22).

##### 3. Capillary velocity

Capillary velocity can be computed from the merger time scale and the final radius of the droplet using an exact relation that was worked out for droplets of a Newtonian fluid in the context of relaxation under a uniaxial deformation [58],

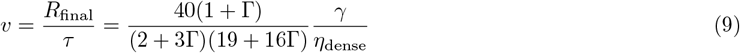

where *τ* is the merger time scale obtained from the fit to the time evolution of shape parameter. Γ = *η*_dilute_*/η*_dense_ is the ratio of the viscosities of the dense and dilute phase. *R*_final_ is the radius of the final droplet and *γ* is the interfacial tension.

The prefactor is determined by the ratio of the viscosity of the dense and dilute phase. The viscosity of the dense and dilute phases have been measured using a rheometer and is presented in the main text in Fig. 4b and Supplement S2 I.

**FIG. S4.**
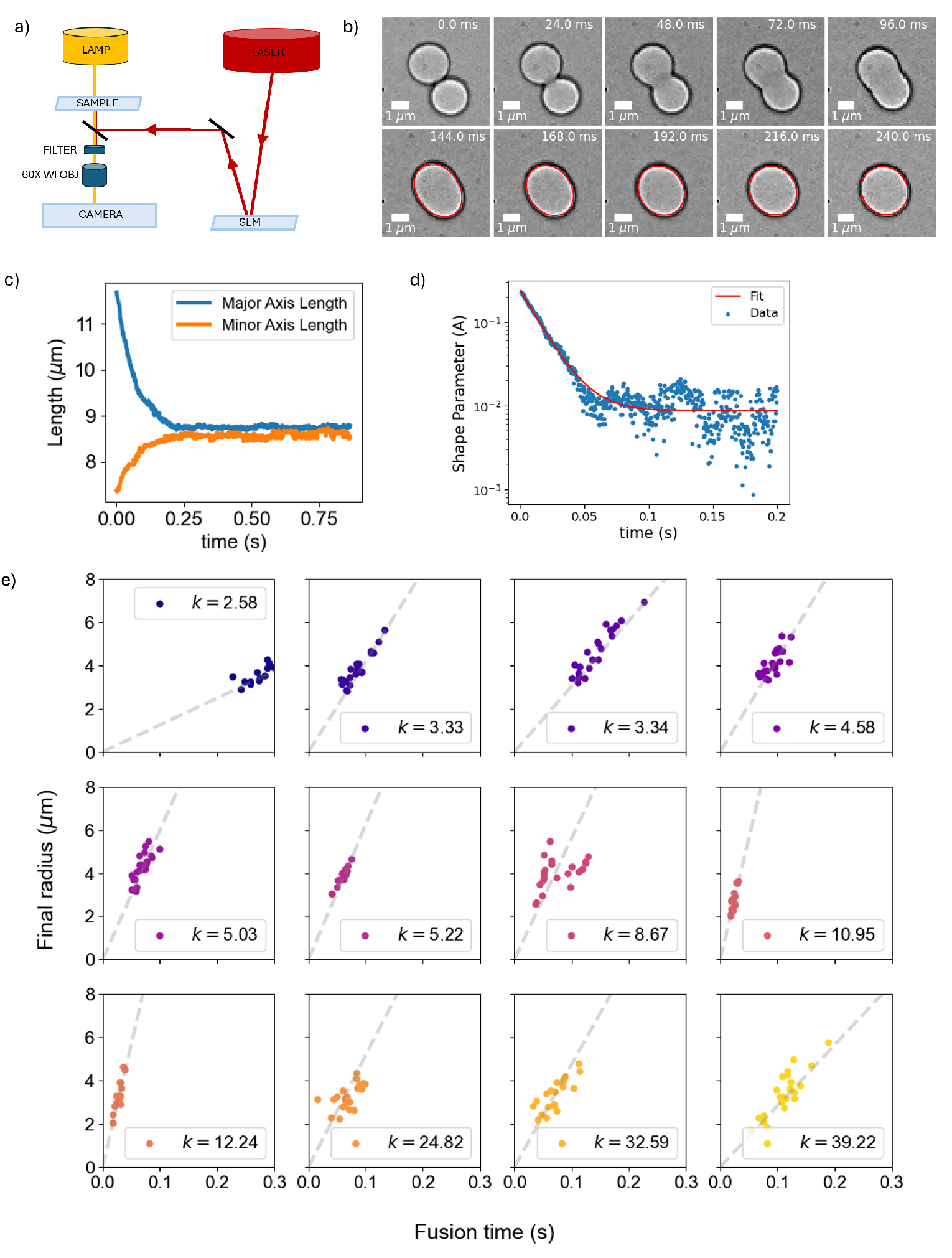
Measurement of capillary velocity. a) Holographic optical trap setup. b) This panel shows a representative droplet fusion event for *k* = 5.22. Two droplets are brought into close proximity to one another and they fuse by randomly colliding with each other due to brownian motion. The lower panel demonstrates the tracking of the droplet as it relaxes from an ellipsoid to a sphere (the line shown in red is the fit to ellipse). c) Shows the major and minor axes lengths obtained from the image analysis of the above merger. d) This shows the shape parameter curves for one droplet mergers at one composition taken at different laser powers. e) Final radius of the droplet plotted against the fusion time. The linear fit is the capillary velocity.

##### 4. Effect of Laser power on capillary velocity

As the optical trap is on during the merging event, we need to ensure that the optical trap itself is not affecting the dynamics of the merger event, for every composition we take between 3-6 droplet mergers at four different laser powers. If we do not see any systematic trend in any of this data, then we can safely conclude that the laser power does not affect the measurement. Fig. S5 shows the capillary velocity for different laser powers for all the compositions of BSA/PEG shown in the main text. We do not see an obvious trend in the data with respect to laser power.

**FIG. S5.**
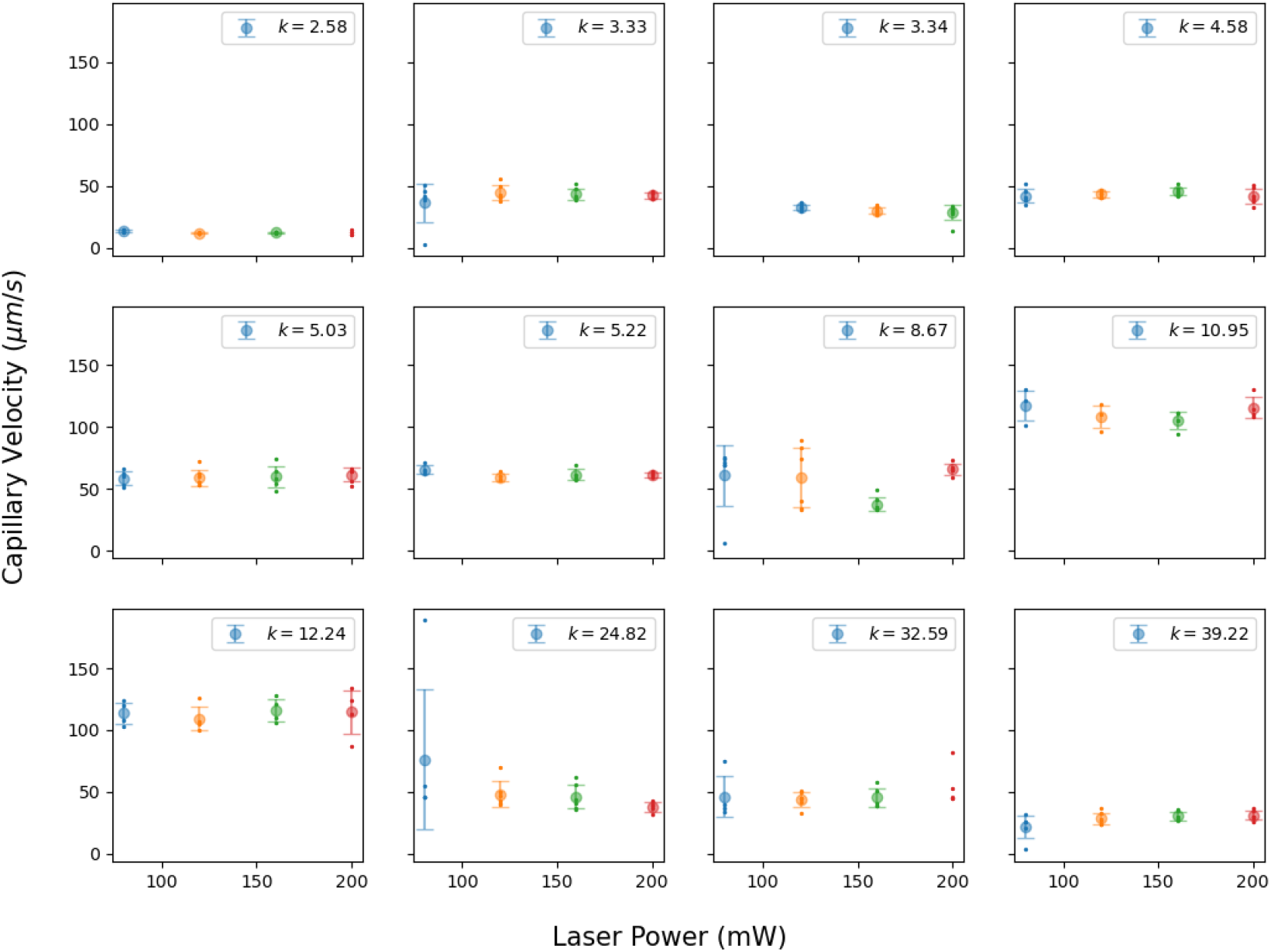
Effect of laser power on the capillary velocity for the range of partition coefficients measured.

#### I. Rheology measurement

The rheology of the dense phase was measured on a TA Instruments DHR3 Rheometer with a 20mm Parallel plate geometry and gap of 500*µ*m. This geometry was chosen so that very small volumes ( ∼50*µ*L) of the dense phase could be used for each measurement. Over 3 minutes, we performed a linear ramp in the strain rate from 0.01 to 1 1/s followed by another 3 minutes of a logarithmic ramp from 10 to 5000 1/s. For these measurements, there was no humidity control and so there will also be a slight amount of evaporation over the 6 mins of measurement time. The temperature was kept fixed at 25^°^C. The raw curves are shown in Fig. S6a.

We note a very large shear thinning effect in the viscosity data starting at 10^8^ mPa · s at 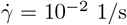 to around 10^3^ − 10^5^mPa · s for 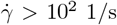. This huge viscosity at low shear rates has been noticed before in BSA solutions and is attributed to a thin film formed at the liquid-air interface [99]. They argue that the high shear rate viscosity is more representative of the bulk because the interface is destroyed at the high shear rates. In order to confirm this hypothesis, we redid this measurement on an AntonPaar TwinDrive Rheometer with a double-gap geometry (DG26.7/Q1-SN52707). This uses a lot of volume of the dense phase ( ∼4mL) and minimizes the interface to air. Logarithmically spaced points were taken at 23C upto 1000*s*^−1^ with the condition of constant value for 4s. This is shown in the red curve. A humidity box was also added to control for evaporation. Notably, we do not see the dramatic shear thinning effect of the parallel plate geometry. The high shear viscosity values are consistent with the high shear viscosity measured in the parallel plate. There is a modest amount of shear thinning from 220 to 160 mPa · s in the dense phase which we will ignore. Therefore, we use the high shear viscosity (averaged from 10^2^ to 10^3^ mPa · s) from the parallel plate geometry as the viscosity of the droplet phase.

For the dilute phase viscosity, we use the same double-gap geometry on the AntonPaar TwinDrive Rheometer with 4mL sample volume. The viscosity was measured for logarithmically spaced shear stress values from 0.001 Pa to 1 Pa and the viscosity was measured with the condition of constant value for 4s. We report the viscosity of the dilute phase at strain rate, 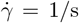 in the main text. The strain rate obtained from the relaxation timescale of droplet fusion is between 1 − 101/s

**FIG. S6.**
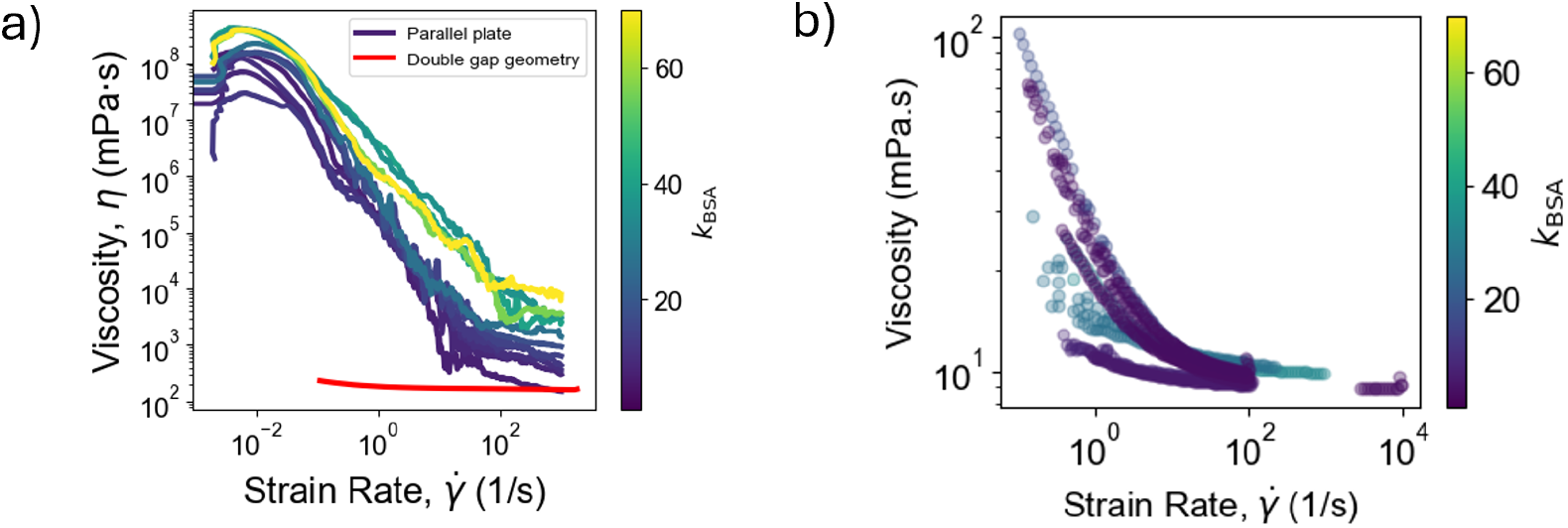
Rheology data. a) Dense phase rheology measured using the parallel plate geometry (partition coefficient indicated by the color bar). Double gap geometry at *k* = 8.8 (red) shows very mild shear thinning behavior and agrees with the parallel plate data at high shear rates. b) Dilute phase rheology data shows very little change in viscosity across the range of partition coefficients measured, roughly 10 times that of water.

#### J. Small Angle X-ray Scattering

SAXS data for the dilute phase and low-viscosity dense phase of the BSA/PEG system were taken using the Cornell High Energy Synchrotron Source (CHESS) with 1.1^Å^ x-rays (11.3 keV). The sample was exposed for 1s and averaged over 10 acquisitions. About 70 *µ*L of the sample was placed in a standard Bio-SAXS quartz flow-through capillary and the sample-detector distance was about 1.8m. This method does not work for very viscous samples. For very viscous samples measurements of pure dense phase were performed at the BioXolver (2.0, Xenocs, Grenoble, France) with 1.54^Å^ x-rays (8keV), typical detector-sample distance of 1.625 m, exposure times of 120 s and averaged over 15 acquisitions. Schematic of the small x-ray scattering experiment is shown in Fig. S7a. The intensity of the x-rays were adjusted so as to not damage BSA protein. Damage of the protein was visualized by the uptick in the curve at low-*q* in the guinier plot, which corresponds to protein aggregation.

##### 1. SAXS structure factor

Curves were analyzed at the beamline using RAW software and using their python API [65]. Form factor *F* (*q*) measurements were taken at a low enough concentration of BSA (10g/L). Fig. S7b shows the form factor measured at CHESS and the in-house Bioxolver. They are in good agreement with each other. Backgrounds for each sample are prepared with the same concentration of PEG, KP Buffer and KCl as the sample. Fig. S7c shows the raw scattering curves along with their corresponding backgrounds. For small concentrations of BSA, there is good agreement at high q between the sample and the background scattering curves, as expected. For very high concentrations of BSA, *e*.*g*, in the dense phase measurements and some dilute phase measurements, the background is well below the scattering curve.

Noting that the high-*q* region is not really changing significantly, we scale the background-subtracted curves so they are sitting on top of each other at high-*q*. Then to obtain the structure factor, we divide out the form factor from the raw intensity profile.

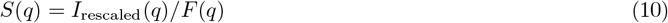

When we align the high-*q* regions of the raw scattering curve with the form factor, we automatically account for the effect of concentration of the protein which introduces an overall multiplicative constant to the curves. We find that this works slightly better than using the measured protein concentrations.

**FIG. S7.**
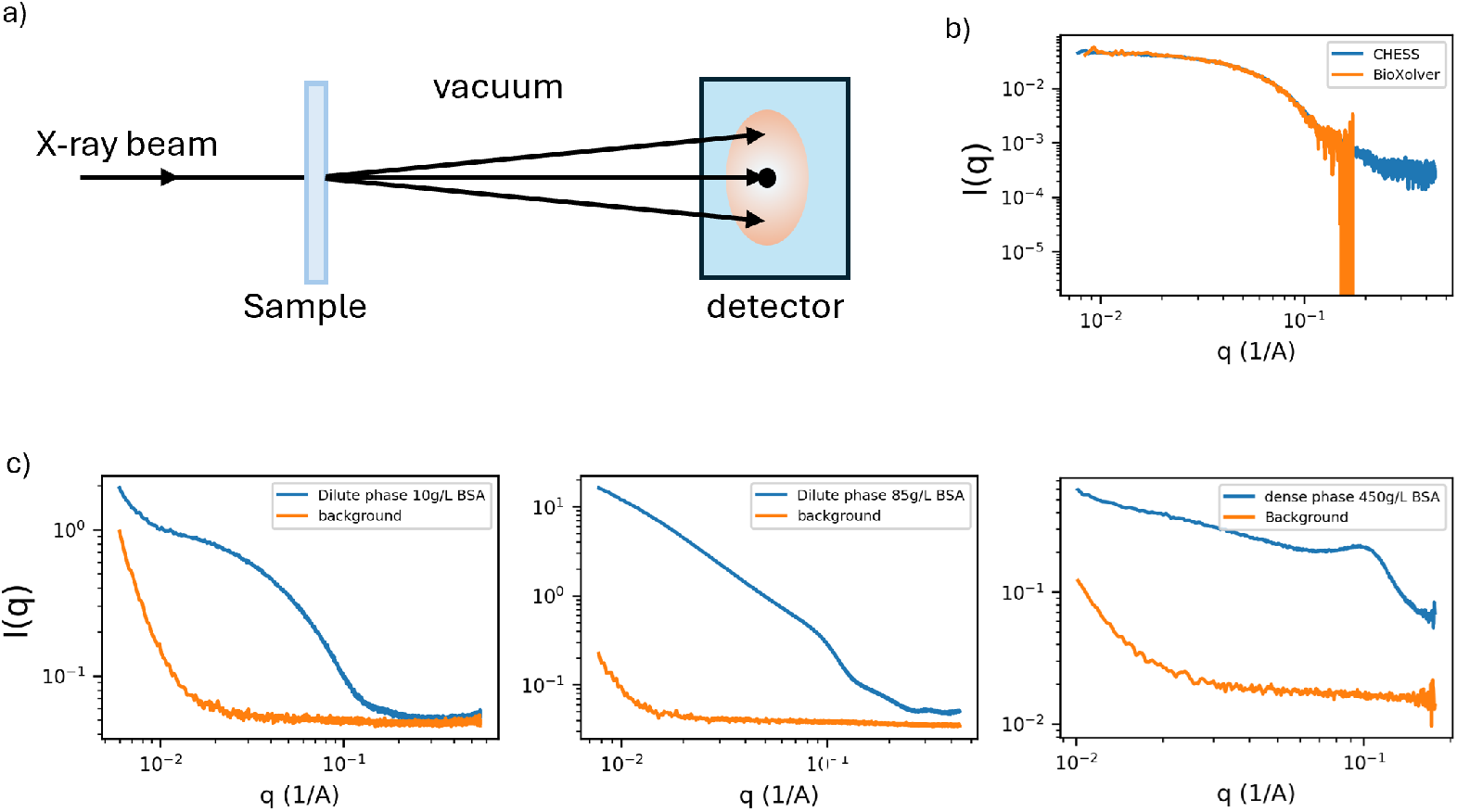
SAXS scattering intensities and backgrounds. a) Schematic of Small angle x-ray scattering. b) Form factor measurements (10g/L BSA) using CHESS and BioXolver. They are in close agreement. c) Scattering from the sample doesn’t match its corresponding background at high-*q* when the protein concentration is high. The conclusions in this paper depend on the structure factor at low-*q* which is barely affected by the exact background subtraction at high-*q*s.

##### 2. Fitting

The curves are fit to a generalized Ornstein-Zernike form [66]:

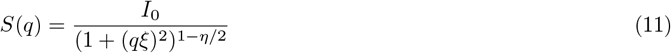

where the fitting parameters are the low *q* plateau, *I*_0_, the correlation length, *ξ*, and the exponent, *η*. This allows us to extract osmotic compression modulus (*K*) from *I*_0_ (see next section) and correlation length *ξ* from the fits. All the structure factors for the dense and dilute phases along with their fits are shown in Fig. S8a-b. The values of the fit parameter *η* and its error is also shown in Fig. S8c.

##### 3. Obtaining osmotic compression moduli

The fit values of *I*_0_ can be converted into an osmotic compression modulus using the following form [64]:

**FIG. S8.**
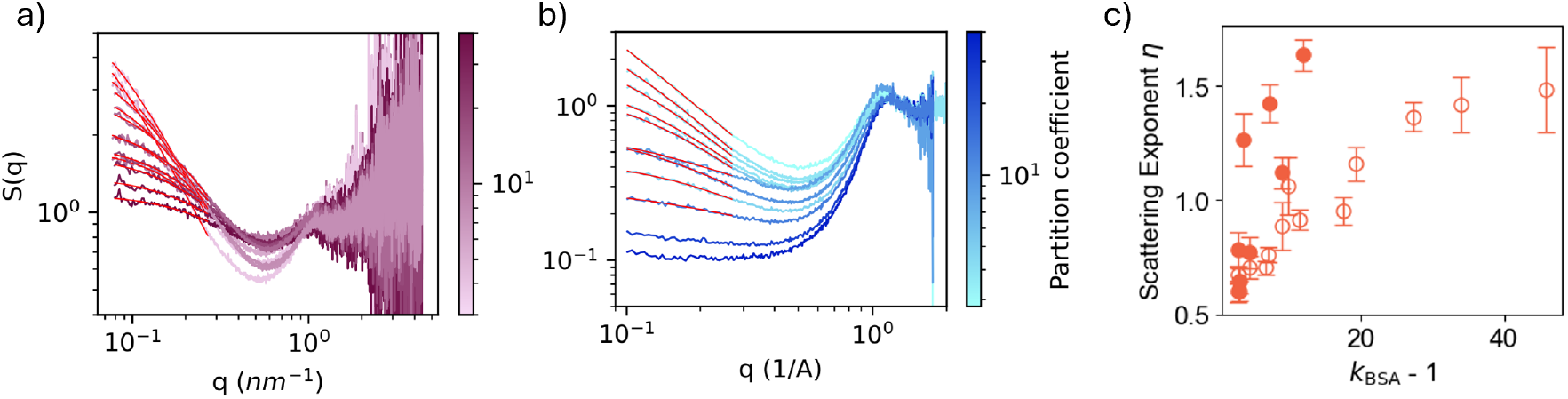
Raw structure factors with fits. a) All the structure factor curves obtained for the dilute phase. The red line through the data is the fit to the generalized Ornstein Zernike function. The color bar shows the range of partition coefficients. b) All the structure factor curves obtained for the dense phase. The red line through the data is the fit to the generalized Ornstein Zernike function. The color bar shows the range of partition coefficients. c) fit values for the exponent *η*. The other fit values are shown in the main text.

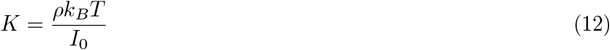

This formula only works sufficiently far from the critical point with small fluctuations. However, we have used this for the whole range of our data.

### K. Estimating volume fractions

We use the form factor of BSA (10g/L BSA in 50g/L PEG, 100mM KP buffer, 200mM KCl) to measure the radius of gyration of BSA in the presence of PEG. We chose the concentration of PEG to be 50g/L as it is a typical concentration of PEG in the dense phase and we do not notice a strong dependence on BSA radius with PEG concentration that will affect the following calculations significantly. The form factor is fit using Guinier analysis to get the radius of gyration of the BSA molecule [100]. The radius of gyration of BSA is measured to be 3.26nm. Assuming a near spherical shape of BSA, we compute the volume fraction of BSA from the number concentration by multiplying the volume of a BSA molecule.

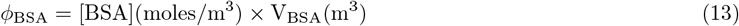

The volume fraction of BSA in the dense phase as a function of the partition coefficient is shown in Fig. S9b. The volume fraction of BSA is approaching the hard-sphere close packing fraction of 0.74.

To convert PEG concentrations into volume fractions, we use the literature value of the hydrodynamic radius of PEG(4k) - 1.65 nm [101] and use the number density of PEG to get the volume fraction.

Using the volume fractions of BSA and PEG we can find the volume fraction of water in the same way as:

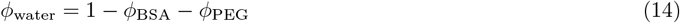

We can now construct the ternary phase diagram (Fig. S9c) and estimate a partition coefficient for water (Fig. S9d).

### L. Fit to 3D Ising model

For the BSA/PEG system, the phase boundary looks nothing like the FUS-urea-water phase boundary since this is a mixture of 2 macromolecules (BSA and PEG) and one small molecule (water). In order to use the law of coexisting densities, we need to generalize the equations 38 and 39 to accommodate multi component systems.

For any component *i* (*i* ∈ [1, *N* )) in an *N* -component mixture, the dilute and dense phase concentrations for that component is given by

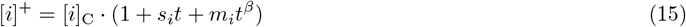

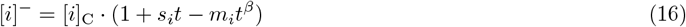

where +*/*− represents dense/dilute phase respectively.

**FIG. S9.**
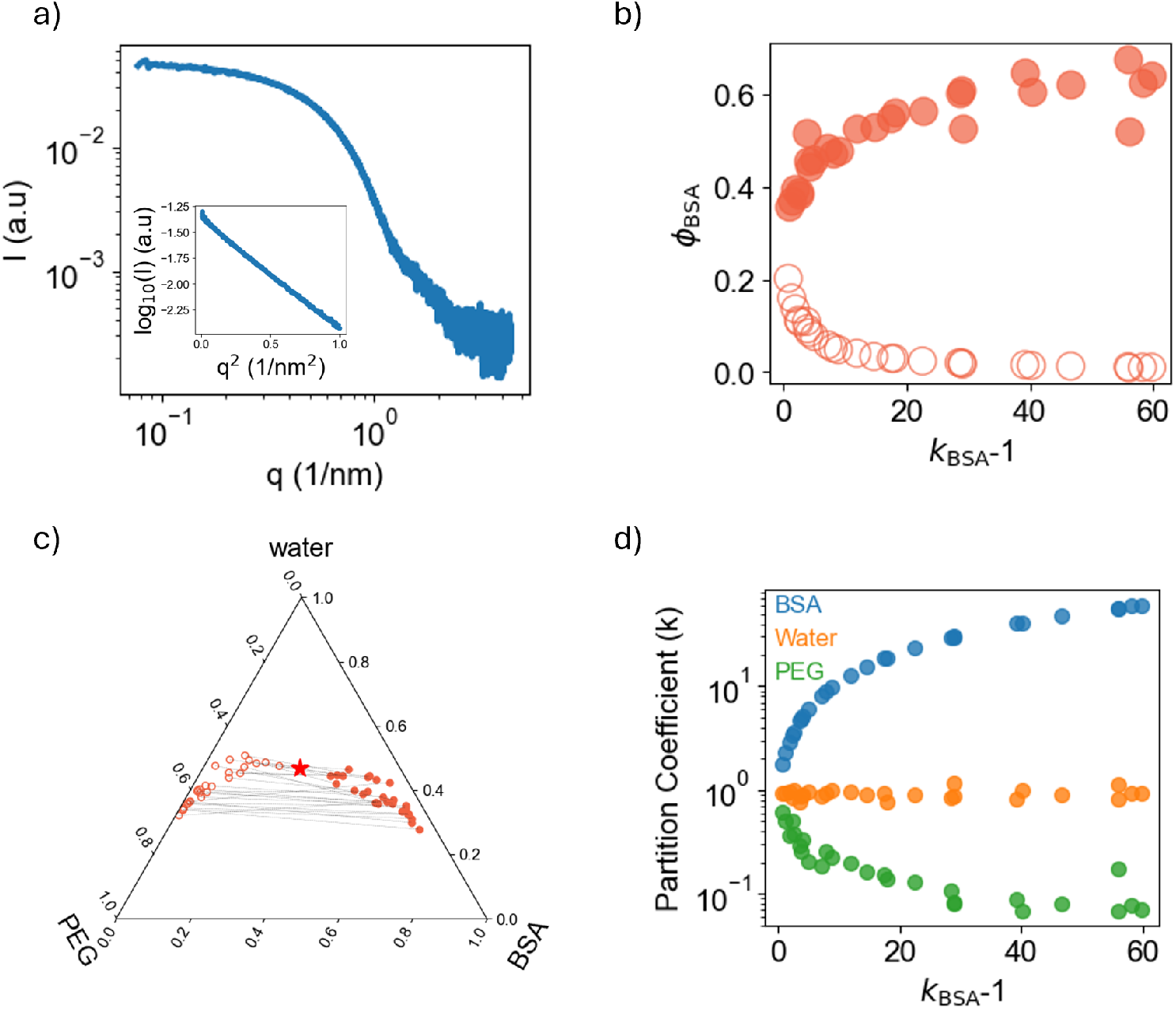
Ternary phase diagram of BSA/PEG system and partition coefficients of individual molecules. a) Form factor of BSA - the scattering from a solution containing 10g/L BSA and 50g/L PEG is taken as the form factor for the BSA in a solution of PEG. Inset shows the Guinier plot of the form factor which is fit to obtain the hydrodynamic radius of BSA. Volume fraction of BSA as a function of partition coefficient computed using the hydrodynamic radius. c) Ternary phase diagram of BSA, PEG, water system d) Partition coefficients of BSA, PEG and water as a function of partition coefficient of BSA.

Specific to the BSA/PEG example, we have:

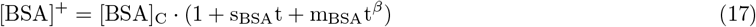

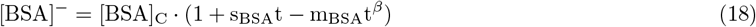

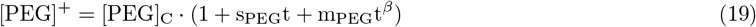

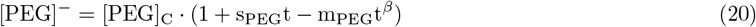

This gives us 6 fitting parameters - [BSA]_C_, [PEG]_C_, s_BSA_, s_PEG_, m_BSA_, m_PEG_.

Again, the scale of *t* is not a fixed and can be changed using the following transformation which leaves the phase boundary invariant,

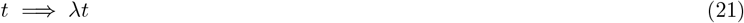

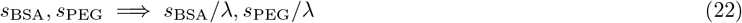

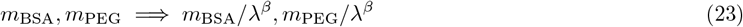

**FIG. S10.**
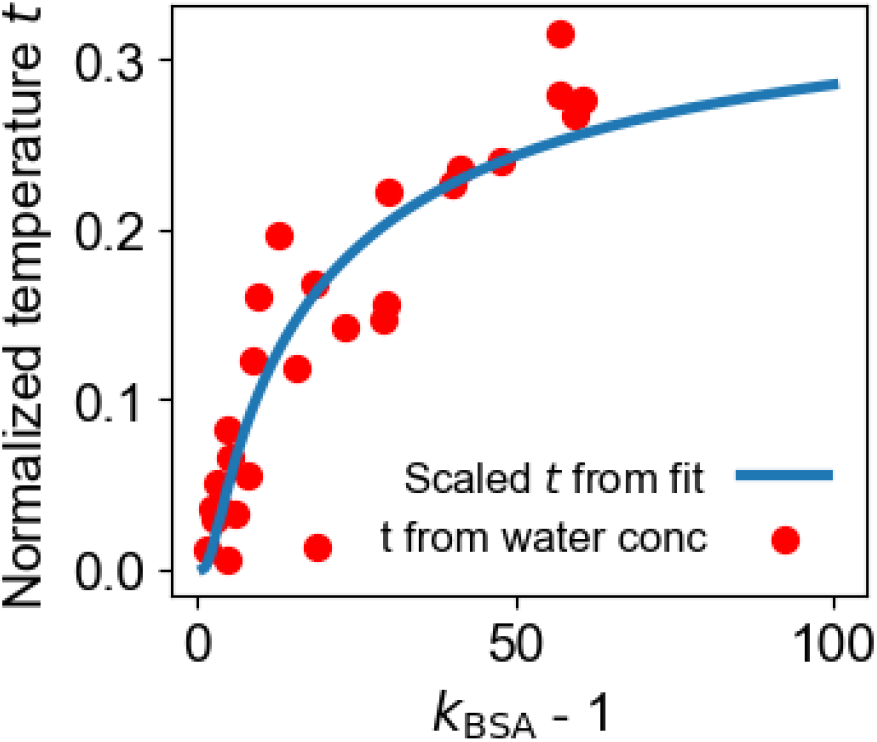
The values of normalized temperature *t* computed using the water concentration from the data is shown by the red circles. The blue line is the value of *t* obtained for a given partition coefficient using the fit to the 3D ising model shown earlier. This has been adjusted by an overall constant scale factor to align with the data.

This invariance leaves only 5 independent fit parameters.

The result of the fit gives the dashed black line shown in the main Fig. 5c) and we can see that it captures the phase boundary across the entire experimental range.

#### 1. Relating t to water concentration

From the ternary phase diagram of BSA,PEG and water, the water partition coefficient is found to be close to one (See Fig. S9d). This implies that tie lines are along lines of constant water concentration in the phase diagram. As discussed in the main text, this enables us to identify the water concentration with the normalized temperature direction.

Analogous to the FUS-urea and water temeprature phase diagram, we can identify the normalized temperature as:

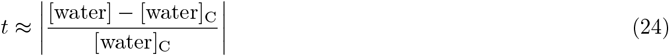

To compare the normalized temperature obtained from the fits to that from our data using the equation above, we plot them on top of each other (see Fig S10). The normalized temperature from the fits have to be corrected by an overall scale (Eq. 21) but we see that they closely follow the *t* computed from water concentration.

### M. Comparison of predicted vs measured exponents for BSA/PEG

The 3D Ising model predicts specific scaling laws for the interfacial tension, correlation length and the compression modulus. In this section, we will compare our experimental findings with theoretical predictions. When fitting our data to obtain power law relations with the normalized temperature, we found that the exponents are very sensitive to the location of the critical point. The location of the critical point is obtained from the fit to the phase diagram. We quantify the range of exponents we would get for a 1% error in the location of the critical point. Larger errors in the location of the critical point produce significantly worse power law fits. We sampled N=5000 different locations of the critical point using a Gaussian distribution with the standard deviation obtained from the fits. While our data appears to be consistent with with the 3D Ising model exponents, the resolution of the experiment does not allow for a tighter estimate of the exponents.

#### 1. Interfacial tension

The interfacial tension is expected to scale as a power law against *t*, with an exponent *µ, i*.*e, γ* ∼ *t*^*µ*^. T The predicted value of *µ* = 1.29 for the 3D lattice gas (equivalent to the 3D Ising model) [77].

We also see a roughly power-law dependence on the *t*-variable.

The interfacial tension scaling with *t* and the distribution of the exponent obtained from the Monte Carlo sampling of the location of the critical point is shown in Fig. S11

**FIG. S11.**
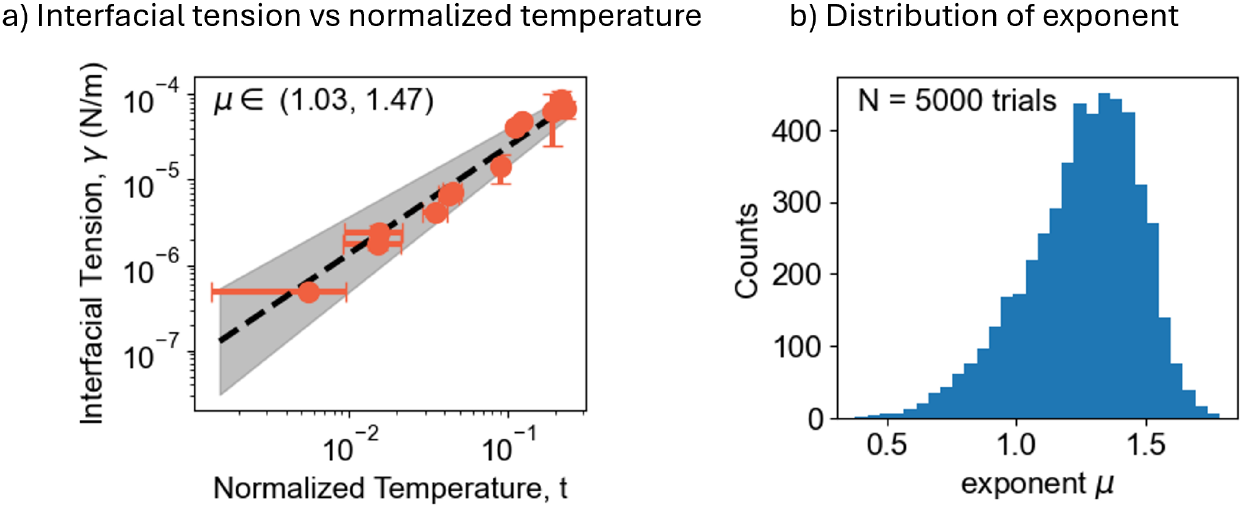
Scaling of interfacial tension. a) Interfacial tension vs normalized temperature. 68% interval of the measured exponent is shown in the top left of the figure (*γ*∼ *t*^*µ*^). The error bars in the normalized *t* direction come from the error bars in the location of the critical point. b) Distribution of the exponent obtained when the location of the critical point is sampled 1% away from the location of the critical point obtained from fitting. The theoretical prediction of exponent *µ* for the 3D Ising model is *µ* = 1.29. [77]

#### 2. Isothermal Compression moduli and correlation length

We perform the same analysis on isothermal compression moduli and correlation length data obtained from the fits to the x-ray scattering as shown in Fig. S12

### S3. FUS-UREA SYSTEM

#### A. Condensate Preparation

For preparation of FUS condensates, FUS stock solution (3500*µ*M FUS, 6M urea, 50mM HEPES, 150mM NaCl) is diluted with FUS buffer (50mM HEPES, 150mM NaCl) by an appropriate dilution factor and thoroughly mixed with a pipette. For susceptibility measurements, additional solutes are mixed into the FUS buffer before adding to the FUS stock. Dilute and dense phases are isolated by centrifuging the sample at 16900g and 23^°^C using the Eppendorf 5418R centrifuge for 10 mins.

#### B. Measurement of FUS concentration

All measurements of the FUS partition coefficient and concentration used fluorescence imaging, on Nikon Ti2 Eclipse microscope equipped with a Yokogawa CSU W1 spinning disk confocal unit, Teledyne Kinetix 22 camera, and Nikon F-LUN laser stack. A 60x water-immersion objective (NA 1.3) was used for all measurements. FUS is labeled with Alexa-Fluor 488 NHS Ester (Succinimidyl Ester) through chemical conjugation. FUS and Alexa-Fluor 488 dye is mixed in 1:1 mole ratio and left undisturbed at room temperature for one hour. The NHS ester of Alexa Fluor 488 forms a covalent bond with free nucleophilic amines on the protein - Lysines and N-terminus of the protein. The excess unreacted free dye is then removed using an Size Exclusion Chromatography (Cytiva HiLoad 26/600 Superdex 200pg) column . The partition coefficient of free dye to FUS condensates is approximately three. We obtain about 16% labeling fraction. Unlabeled protein stock is mixed with the labeled protein at appropriate volumes to achieve 3% labeling fraction.

**FIG. S12.**
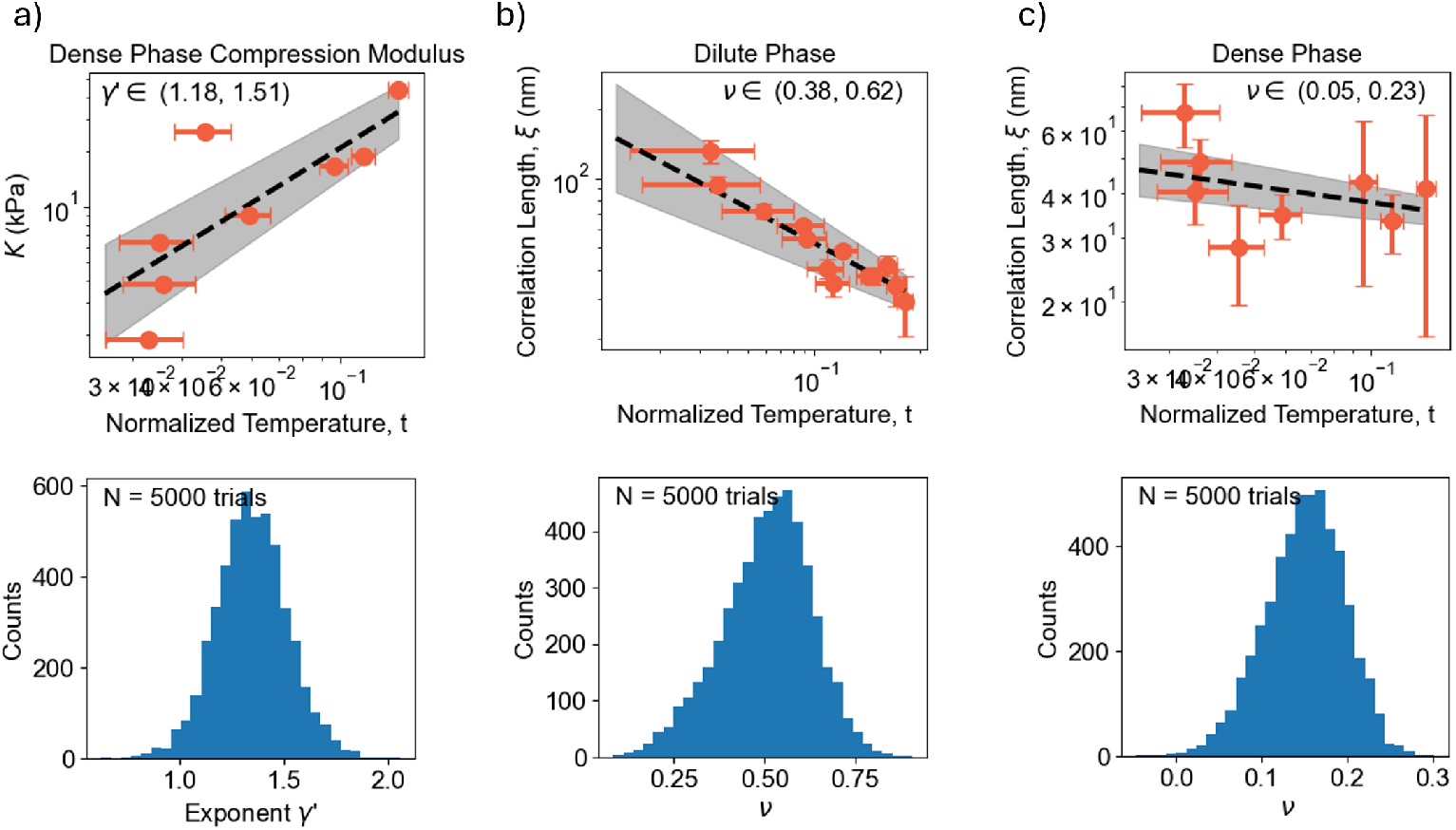
Scaling of Compression moduli and correlation length. a) Fit for power law (top) and the distribution of exponent (bottom) for the compression modulus of the dense phase (*K* ∼ *t*^*γ*^ ). The 68% confidence interval of the exponent is shown on the top right plot and the predicted value is *γ*^*/*^ ≈ 1.31 [78] b) Fit for power law (top) and the distribution of exponent (bottom) for the correlation length of the dilute phase (*ξ* ∼ *t*^−ν^ ). The 68% confidence interval of the exponent is shown on the top right plot and the predicted value is ν ≈ 0.629. c) Fit for power law (top) and the distribution of exponent (bottom) for the correlation length of the dense phase (*ξ* ∼ *t*^−ν^ ). The 68% confidence interval of the exponent is shown on the top right plot and the predicted value is ν ≈ 0.629 [66].

For the calibration, we image homogeneous solutions of the labeled FUS at different concentrations and fit a straight line through it. The dense phase concentration always involves extrapolating this straight line up to three times of the maximum concentration used in the calibration.

We obtain z-stack images of all condensates and identify individual droplets using a thresholding algorithm. We locate the mid plane of each droplet and measure the intensity profile from the center of the droplet up to 2.25 times the radius of the droplet in this plane. The intensity is converted to a concentration using the calibration ladder. Finally we identify the plateau in concentration in the bulk of the droplet as the dense phase concentration and the plateau outside of the droplet as the dilute phase concentration. Between 100 to 1000 droplets are analyzed per sample (see Fig. S13)

#### C. Susceptibility measurements

To measure the dilute phase susceptibility of the condensate with respect to a solute, five phase-separating mixtures whose average composition varies only in the solute concentration, are prepared. The dilute phase of the phase-separated mixture is isolated by centrifuging the sample for 10 mins at 16900g at 23^°^C with the Eppendorf 5418R centrifuge. The protein concentration of the dilute phase is measured using the Nanodrop UV-Vis spectrometer. The dilute phase is diluted with an appropriate buffer (6 M urea, 50 mM HEPES, 150 mM NaCl) so that absorbance at 280 nm lies between 0.05 and 10 A.U. Extinction coefficient for FUS used is 0.0432 *µ*M^−1^cm^−1^. For measurement of FUS susceptibility to urea, arginine, 1,6-hexanediol and proline, the dilute phase concentration of FUS is measured using the absorbance at 280nm. For the case of UMP and ATP, they contribute to the absorption at 280nm. Therefore, we cannot measure dilute phase FUS concentrations by just measuring the 280nm absorbance of the dilute phase. Instead, we use fluorescently labeled FUS molecule (with Alexa 488) and calibrate the absorption of the dye (at 495nm) with the FUS concentration shown in Fig. S14b. Thus, dilute phase FUS concentrations are measured with the 495nm absorbance of the dilute phase using fluorescently labeled FUS. Each dilute phase is measured three times to obtain error bars for the measurement. A straight line is fit between the initial average solute concentration and the corresponding dilute phase protein concentration to obtain the dilute phase susceptibility of the condensate to the solute (fits are shown along with data in Fig S14).

**FIG. S13.**
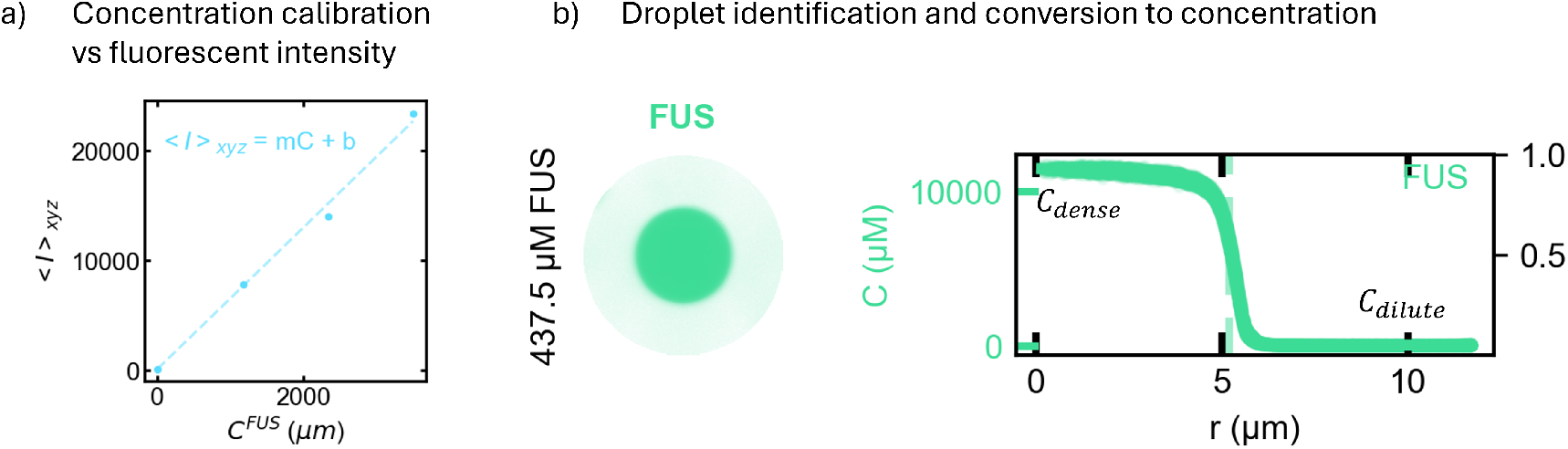
Measurement of concentration using Fluorescence. a) Calibration of fluorescent intensity vs concentration of FUS. b) Using confocal images, we identify droplets in the FOV and measure the intensity of the droplet as a function of its radial distance. The intensity is converted to concentration using the calibration curve and thus we obtain the dense and dilute compositions for each droplet.

#### D. Measurement of FUS partitioning with amino acid mixture and urea

FUS partition coefficient is the ratio the dense and dilute phase concentrations of FUS. It is estimated using fluorescence spectroscopy as described in Supplement S3 B.

##### 1. Composition of 12 amino acid mixture

The stock concentration of the 12 amino acid mixture is shown in table S1.

**TABLE S1.** Composition of amino acid mixture. The name of the amino acid, the target concentration in mM and mg/mL, solubility limits of each amino acid at 23C and the actual concentration of amino acid in stock solution is provided. Solubility limits were found by looking up individual amino acids in PubChem [102]

| Amino acid | Target Conc (mM) | Target Conc (mg/mL) | Solubility Limit (mg/mL) | Actual conc (mg/mL) |
| --- | --- | --- | --- | --- |
| Proline | 240 | 27.6312 | 162 | 27.6312 |
| Glutamic acid | 240 | 39.636 | 8.57 | 8.57 |
| L-Arginine | 240 | 41.808 | 182 | 41.808 |
| Isoleucine | 240 | 31.4808 | 34.4 | 31.4808 |
| Aspartic Acid | 240 | 31.944 | 5.39 | 5.39 |
| Serine | 240 | 25.416 | 425 | 25.416 |
| Glycine | 240 | 18.0168 | 249 | 18.0168 |
| L-Alanine | 240 | 21.384 | 164 | 21.384 |
| Lysine-HCl | 240 | 43.836 | 1000 | 43.836 |
| L-Glutamine | 240 | 35.0736 | 41.3 | 35.0736 |
| L-Threonine | 240 | 28.608 | 97 | 28.608 |
| Leucine | 240 | 31.4808 | 21.5 | 21.5 |
|  |  |  | Total g/L conc of amino acids: | 308.7144 |

While making the stock concentration with a target of 240mM for each amino acid, not all amino acid was completely dissolved due to their limited solubility. The stock solution was centrifuged and the supernatant was extracted. The concentration of a particular amino acid in the supernatant is either the target concentration or the solubility limit of the amino acid. This is also shown in the table above and the final concentration of all amino acids in the supernatant is shown. The supernatant is diluted to various concentrations and added to condensate solution.

**FIG. S14.**
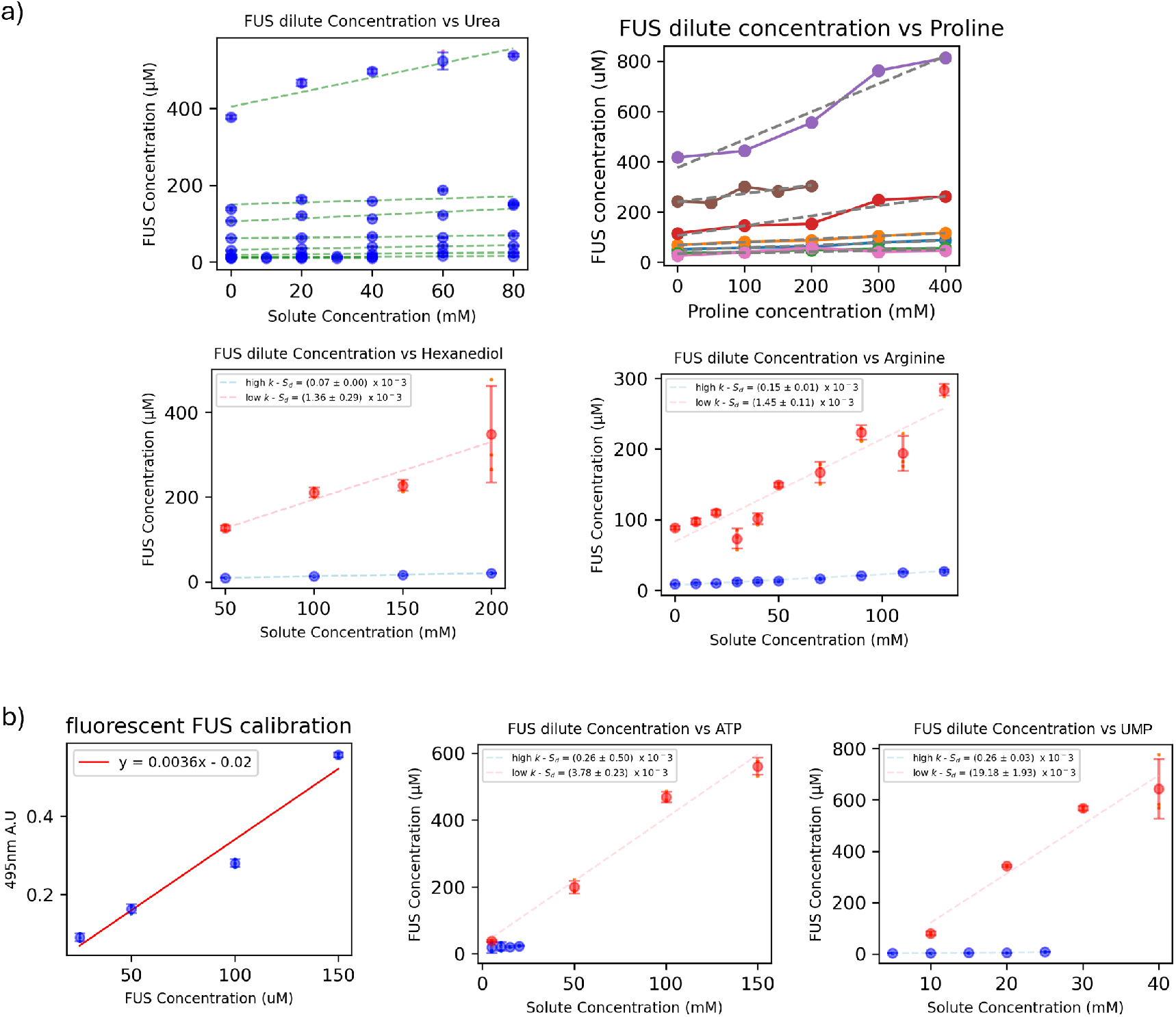
Raw data for susceptibility measurements of FUS to solutes shown in the main text. a) Susceptibility curves for FUS b) Calibration of absorbance at 495nm of fluorescently tagged FUS molecule with FUS concentration and raw susceptibility curves for ATP and UMP.

##### 2. Quenching assays for Urea and amino acid mixture

Urea and free amino acid mixtures are titrated in at different concentrations to free Alexa 488 dye at 10*µ*M concentration. We observe no significant quenching of the dye (Fig. S15)

**FIG. S15.**
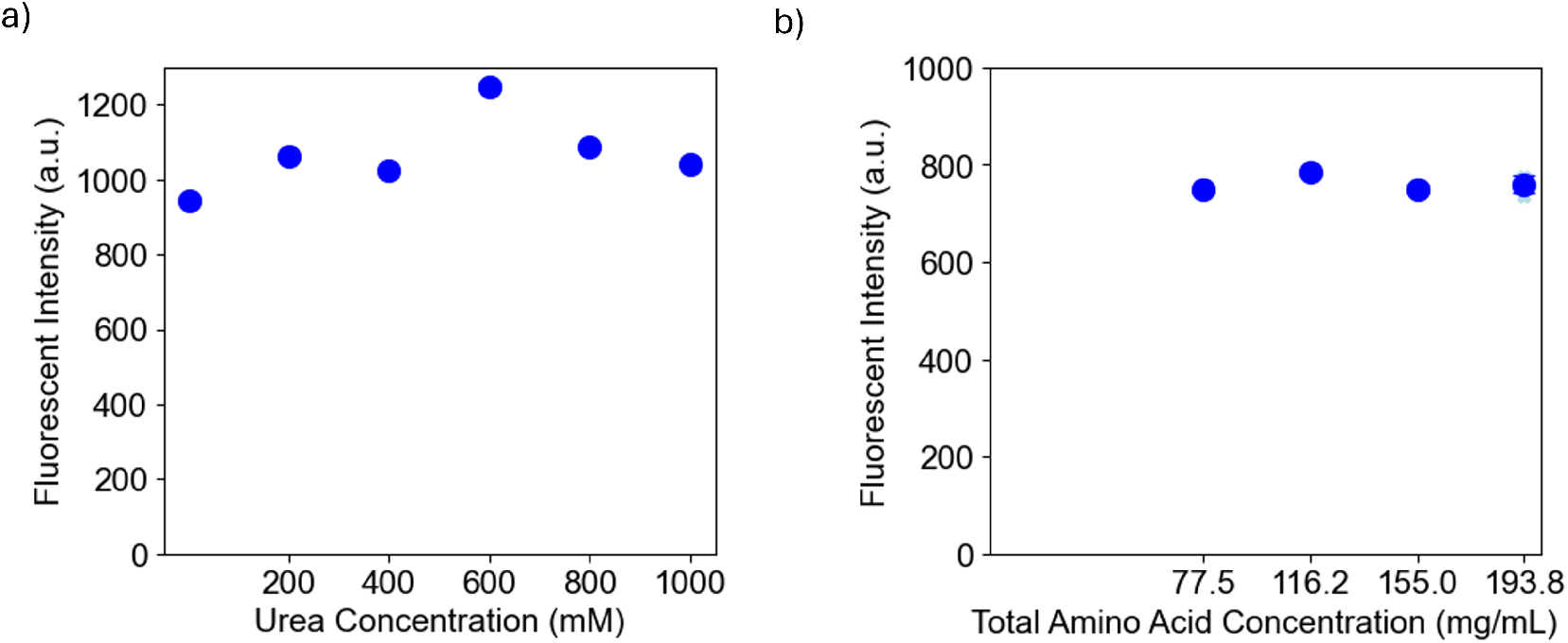
Quenching assays of Alexa 488 fluorophore. a) Free Alexa 488 dye at a constant concentration (10*µ*M) is titrated with up to 1000mM urea and the fluorescent intensity is measured at 20% Laser power. b) Free Alexa 488 dye at a constant concentration (10*µ*M) is titrated with Amino acid mixture between 70 and 200 mg/mL and the fluorescent intensity is measured at 30% Laser power.

##### 3. Corroborating results with UV-Vis spectroscopy

To corroborate our fluorescence results, we measured the concentration of FUS in the dilute phase using UV-Vis spectroscopy. Due to the very small volumes of dense phase, it is not feasible to measure the dense phase concentration using UV-Vis spectroscopy. Both the approaches show a similar trend with an overall offset (See Fig. S16).

**FIG. S16.**
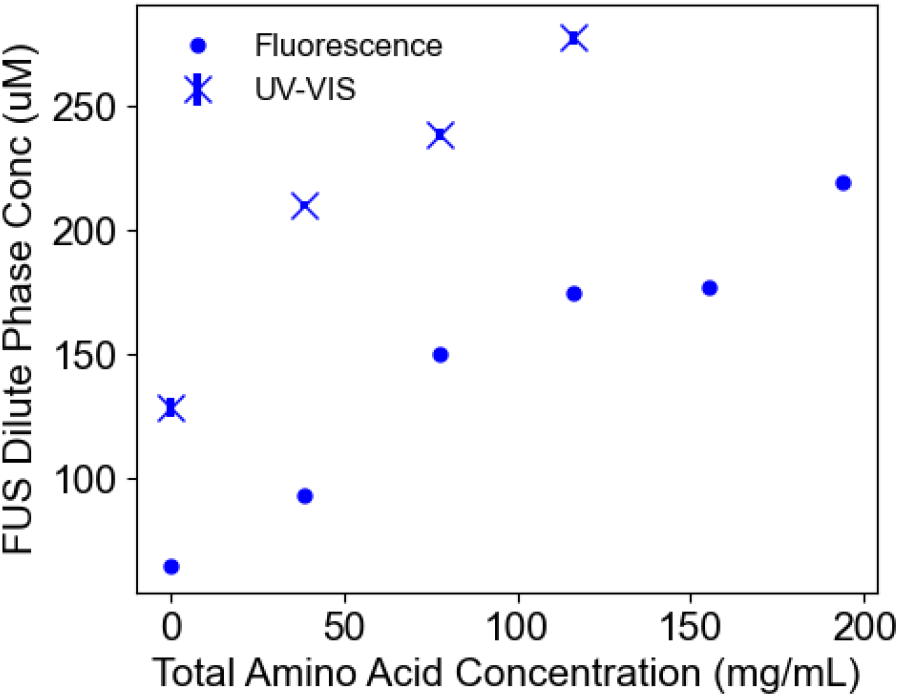
The dilute phase FUS concentration measured using fluorescence (circles) is compared to that measured using UV-Vis spectroscopy (crosses).

#### E. Measurement of urea partitioning: Raman Spectroscopy

##### 1. Data Collection

Measuring urea concentration in FUS condensates is challenging because the volume fraction of the condensates is very low making it hard to isolate them. We used confocal Raman spectroscopy in order to measure the urea concentration inside the FUS condensate. All confocal Raman measurements were performed using the WITec Alpha300R Confocal Raman Microscope with the 532nm laser with 300 l/mm diffraction grating. 50x LWD objective with 0.5 NA was used.

Fig. S17a) shows the optical ray diagram of the experiment. Laser light at 532nm is incident on the droplets and focused into the sample. Light gets back-scattered from the sample and all the light that emerges from the confocal volume is focused into the detector. In order to isolate the inelastic Raman scattering of the back-scattered light, there is a 532nm filter in place before it hits the diffraction grating and the detector. The confocal volume is 1 *µ*m across and 2.5 *µ*m in depth. From the Raman spectra, we can identify different molecular bonds present in the confocal volume as they will have a different wavelength shift based on their energy.

To focus into the droplet, we collect Raman scattering spectra for different heights above the glass slide in 1*µ*m steps. The spectra collected at different heights is shown in Fig. S17c). The peaks observed at different Raman shifts correspond to different molecules present in the confocal volume.

As a rough estimate for the FUS concentration in the confocal volume at each height, we look at the ratio of the intensity of the signal coming predominantly from FUS (C-H stretching at 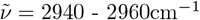) to a signal coming predominantly from water (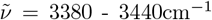) as indicated in Fig. S17c. As the confocal volume sweeps the droplet, this ratio must be a constant. At a certain height, the confocal volume is no longer enclosed within the droplet and the scattering now also contains scattering from the dilute phase. The FUS concentration in the confocal volume therefore decreases eventually plateauing to the dilute phase value. Using this we can determine the height of the confocal volume that is focused inside the droplet. In practice, we do this for one droplet in each sample, and maintain the same height while measuring other droplets in the same sample. In order to test that the 532nm laser is not burning or damaging the protein in the sample, we measured the Raman spectrum at exactly the same spot again and got exactly the same scattering signal as shown in Fig. S17d which proves that the laser does not damage the proteins. To measure the dilute phase, we move to a location without any droplets and move 30*µ*m above the measured droplet height to minimize interference with the glass.

##### 2. Data Analysis

###### Background Subtraction

Raman scattering is also accompanied by a lot of background intensity typically coming from fluorescence of the molecules. The background is assumed to be a smoothly changing lower envelope which captures the fluorescence features. The background is calculated using an asymmetrically reweighted penalized least squares smoothing (ArPLS) algorithm [103] with parameter *κ* = 10^−5^. We find that changing *κ* affects the final urea concentrations measured and have included this while computing error bars.

###### Obtaining spectra of individual components of mixture

The Raman spectra of a solution mixture is the linear sum of the Raman spectra of each molecular species present in the mixture proportional to its concentration. The FUS dense phase is a mixture of urea, FUS, water, HEPES and NaCl. Since the concentration of HEPES and NaCl relative to water is constant across all samples (50mM HEPES and 150mM NaCl), we can think of this as one entity henceforth referred to as the FUS buffer. There is also a mixing of the glass signal in the dense phase due to the proximity of the glass slide to the droplet. Therefore we will include that also in the scattering spectrum.

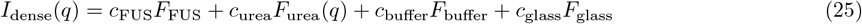

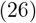

where *c*_x_ is proportional to the numbers of x molecules and *F*_x_ is its Raman spectrum (x = FUS, urea, buffer, glass).

To obtain individual urea and buffer Raman spectra, we first run a calibration ladder of different urea concentrations in the FUS buffer. We notice that the N-H stretching peaks of urea affect the O-H stretching of the water also. Therefore, in order to correctly extract the urea and FUS buffer signals from this data, we invert Eq. 26 for the calibration spectra as shown in Eq. 27 (Our mixture only consists of urea and buffer).

**FIG. S17.**
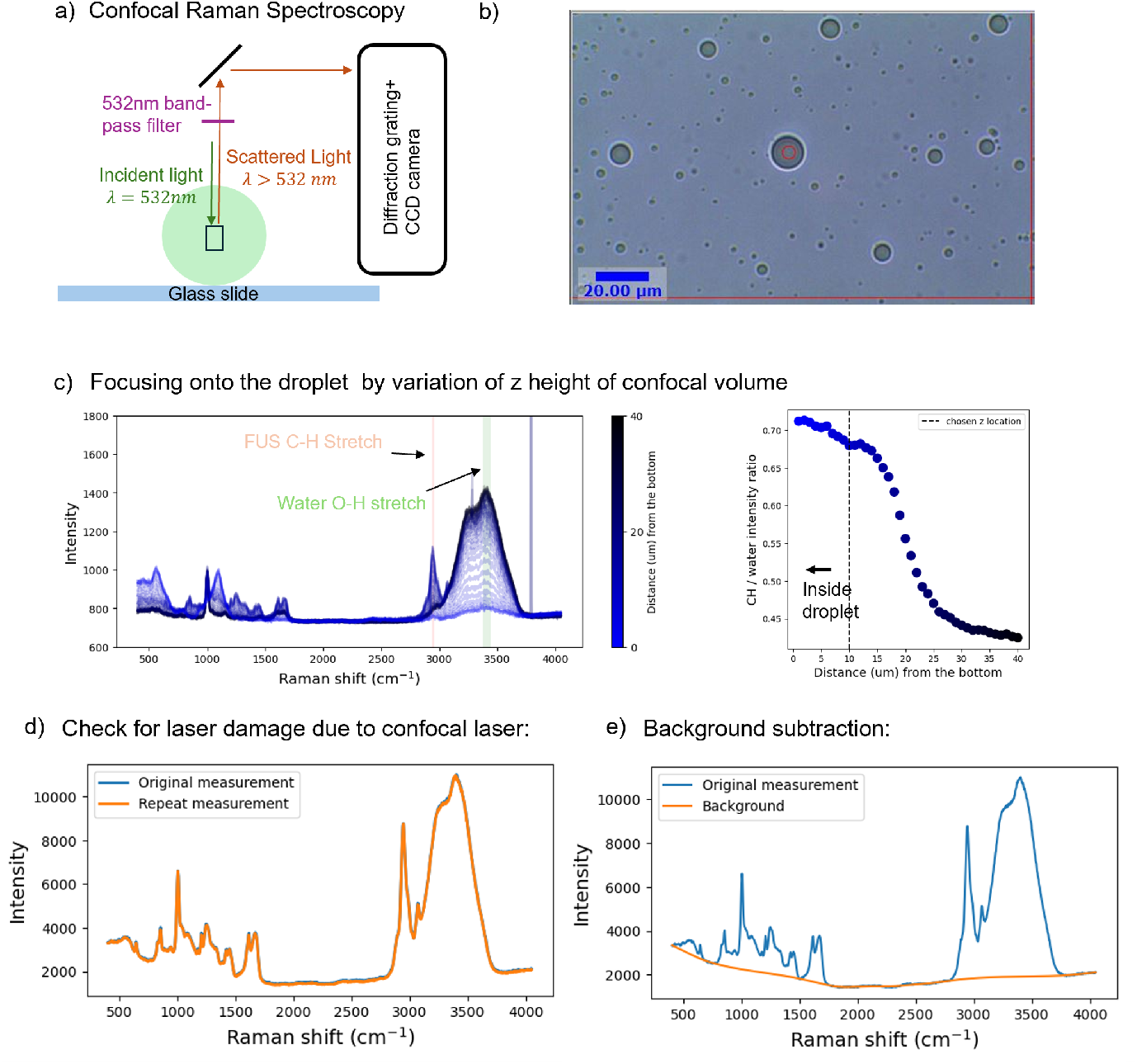
Raman data collection. a) Schematic of the Confocal Raman setup. b) Typical image of condensates viewed under the bright-field setting of confocal Raman microscope. The laser is focused onto a point inside the red circle at the center of the droplet. The width of the laser spot is smaller than the red circle. c) Spectra from within the confocal volume as a function of height above the coverslip from a z-scan (left). Also shown are the C-H stretching peaks predominantly coming from FUS and O-H stretching peaks predominantly from water. To identify the inside of the droplet, we compute the ratio under the small range of Raman shifts shown in the figure. The ratio as a function of distance from the glass slide is shown on the right. d) We irradiate the condensate at the same spot to measure any changes in the Raman spectrum. e) typical background computed for the Raman spectrum.

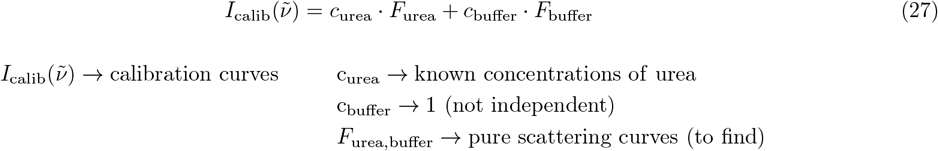

The LHS of the equation are the measured curves of the calibration ladder. We know the concentration of urea that was used in the experiment. Concentration of buffer is not an independent quantity and only affects the overall normalization of the curve. Therefore, we set the concentration to be 1 for all the curves. The closest pure buffer and urea curves (*F*_urea_, *F*_buffer_) that satisfy the above equation is fit using a least squares procedure (lstsq function in Numpy linalg library). The calibration curves and the corresponding to urea and buffer curves obtained from the lstsq fit is shown in Fig. S18 b-c.

The spectrum of glass was measured by focusing the confocal volume purely inside the glass slide (Fig. S18e).

In order to obtain the Raman spectrum of FUS, the stock concentration of FUS (3500 *µ*M) dissolved in 6M urea was not at a high enough concentration to resolve the finger print region (500-2000 cm^−^1) of the protein. Instead, we dilute the stock by 200 times using the FUS buffer until there is a total of 30mM urea in the solution. The FUS condenses into droplets at this urea concentration, but the urea concentration within the droplets is not high enough to be detected. The corresponding signal thus consists mostly of FUS and FUS Buffer. To obtain the contribution of FUS to the spectrum, we subtract out the FUS Buffer spectrum to obtain the curve Fig. S18b.

**FIG. S18.**
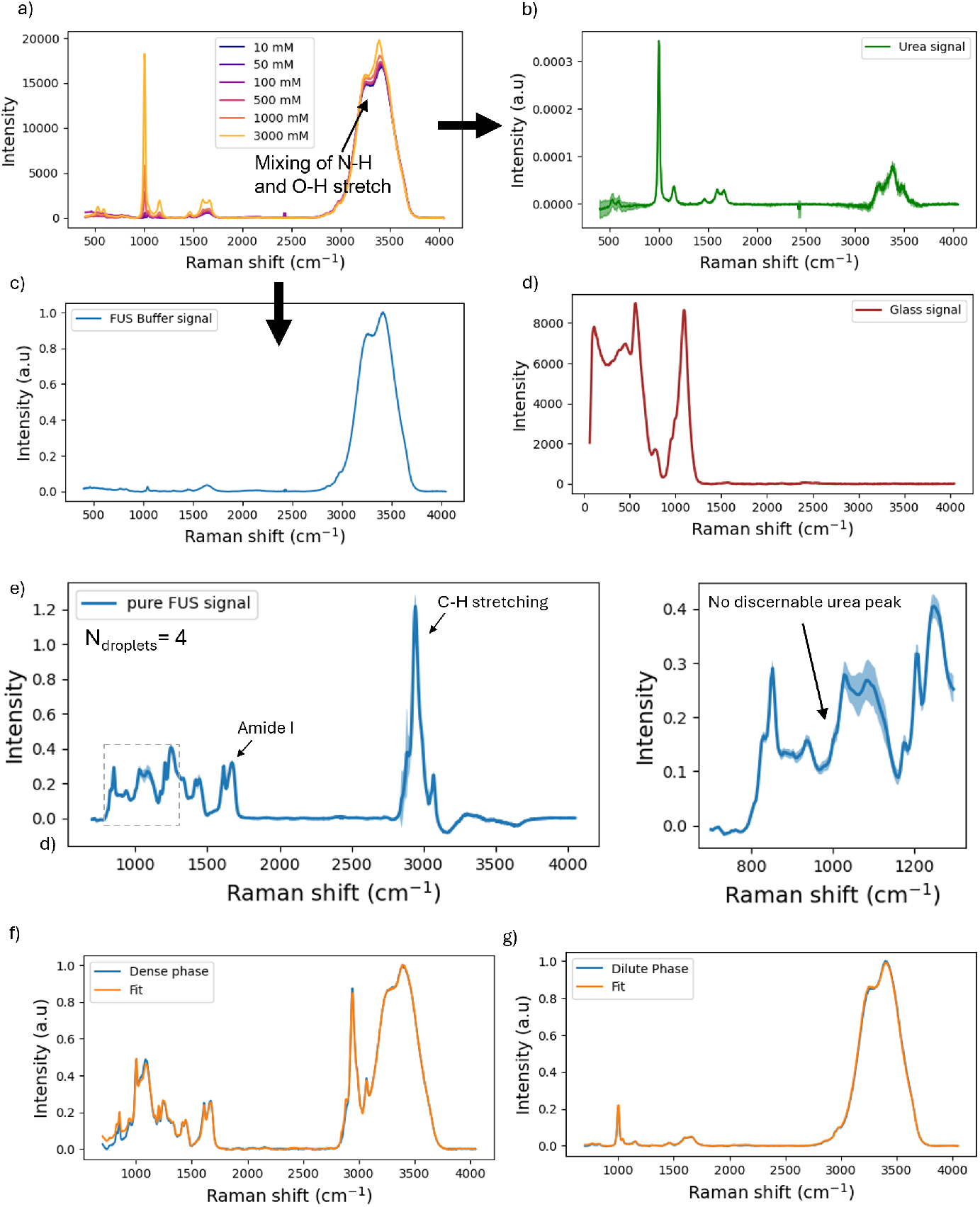
Raman data fitting. a) Raman spectra for mixtures of urea and FUS buffer with varying concentration of urea. There is clear mixing of N-H and O-H stretching of the urea and water around 3000 - 3500 *cm*^−1^ Raman shift. b-c) Urea and buffer spectra extracted from least squares fitting of the calibration curves in (a). d) Pure glass signal measured by focusing the confocal volume into the glass slide. e) The scattering from condensates in the presence of very less urea (30mM) is approximated as the scattering from pure FUS protein (shown on left). The buffer signal is subtracted out to obtain scattering from just FUS. The urea peak is barely visible at this concentration. f-g) Fitting of the dense and dilute phase spectrum using the above measured pure signals.

###### Fitting the spectra dilute and dense phase mixtures to extract concentrations

Now that we have the pure buffer, urea, FUS and glass signals, we can now fit the full background-subtracted spectra of the dense and dilute phase using Eq.26 with the spectra of individual components.

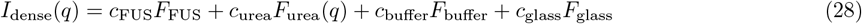

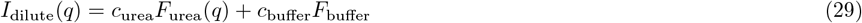

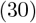

Where the concentrations are fit parameters. The dilute phase signal does not contain a measurable FUS signal and so we do not include it in the fit. Using least squares fit, we can find what are the concentration values that best approximate the dense and dilute phase scattering of the sample. The fits are shown in S18 f-g. Again, there is an overall normalization factor and the concentration of urea is the ratio of *c*_urea_*/c*_buffer_

###### Interpolation of urea partition coefficient

From the concentration of urea in the dense and dilute phase, we can plot the partition coefficient of urea *k*_urea_ as a function of the dilute phase urea concentration. We measure the urea concentration to be between 0.5 − 1.1 for the range of urea concentrations measured. For the measurements, it is hard to distinguish deviations away from one within the error bars. Therefore, when plotting phase diagrams, we assume the urea partitioning to be equal to one.

**FIG. S19.**
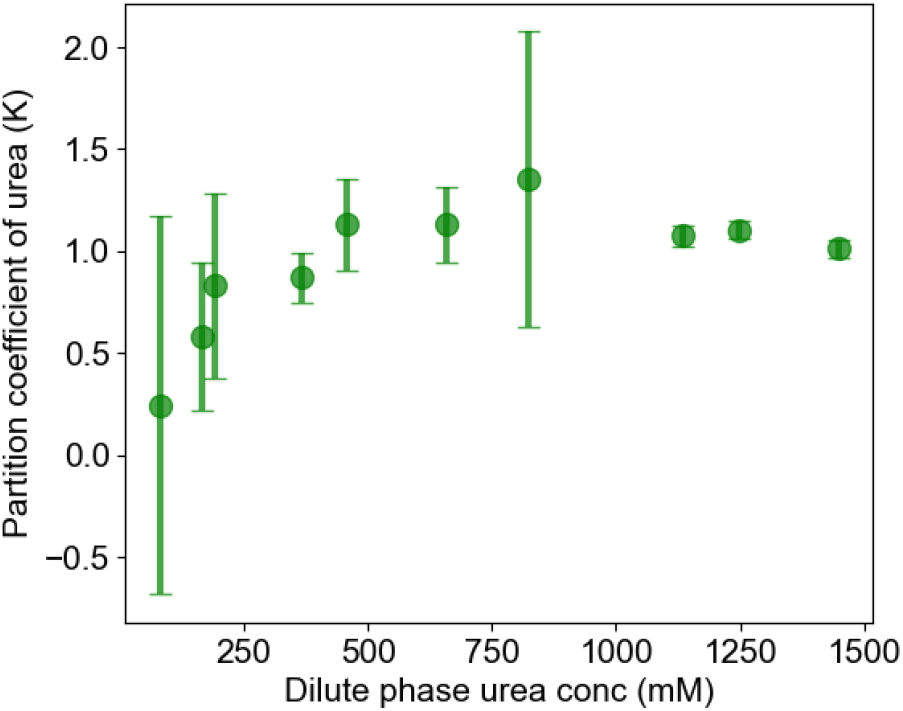
Partition coefficient of urea measured using Raman spectroscopy vs the dilute phase concentration of urea.

#### F. Fit to 3D Ising model

In analogy to the liquid-vapor fit, the coexisting dilute and dense phase of FUS condensates can be fit using the 3D Ising model phase diagram. As mentioned in the main text, urea in the FUS-urea system acts analogous to temperature in the temperature-water system. We can write down the normalized temperature using the urea concentration as,

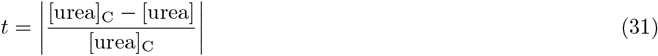

where [urea]_C_ is the concentration of urea at the critical point. We use the same fitting form as for the liquid-vapor phase diagram (Eq. 38, 39) but this time with a newly identified urea-based normalized temperature variable.

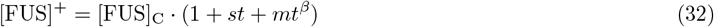

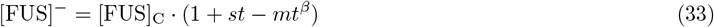

The fitting parameters are now {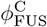, *s, m*, [urea]_C_} and are found using a least squares minimization procedure.

#### G. Flory-Huggins fit

To compare the FUS-urea binodal to those predicted from the Flory-Huggins theory [81], we must first convert FUS and urea concentrations from molar units to volume fractions. FUS concentration (in *µ*M) is converted to mg*/*mL units using the molecular weight of FUS (23.6kDa) and then converted to volume fractions using the density of the amino acid, following [104].

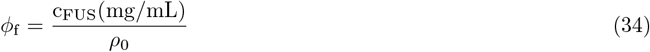

Where *ρ*_0_ is the density of an amino acid given by *ρ*_0_ = 1310.16 mg*/*mL.

In order to convert urea concentrations from mM units to volume fraction, we need an approximate volume of the urea molecule (*v*_urea_), approximately 0.0796 nm^3^ [105]. Then, the volume fraction of urea is given by:

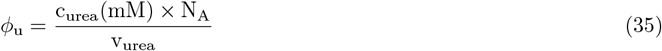

Where *N*_*A*_ is the Avogadro’s number.

Using the volume fraction of urea and FUS, we can find the volume fraction of water as:

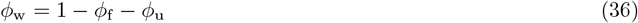

The Flory-Huggins theory at a fixed temperature for the FUS-Urea-water system is given by,

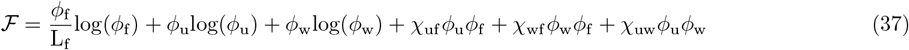

where L_f_ is the size of the FUS molecule relative to water and *χ*_uf_, *χ*_uw_, *χ*_wf_ are pairwise interactions between the molecules.

The composition of the coexisting phases were computed from this free energy using the convex hull algorithm [83]. The free energy was sampled on a 600× 600 grid (360,000 points). *ϕ*_*u*_ was linearly spaced from 0 to 0.15, while *ϕ*_*f*_ was logarithmically spaced from 0.0002 to 0.8. *ϕ*_*w*_ was computed as *ϕ*_*w*_ = 1 − *ϕ*_*f*_ − *ϕ*_*u*_. The convex hull was computed of this free energy and the number of phases and the composition of coexisting phases were found. The phase boundary was fit by hand and the parameters best representing the phase boundary were:

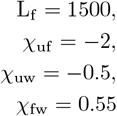

### S4. BIK1-NACL SYSTEM

#### A. Condensate preparation

For preparation of Bik1 condensates, stock solutions of Bik1 (731 *µ*M Bik1, 500mM NaCl, 20mM Tris) is diluted with Bik1 buffer (20mM Tris) at appropriate dilution factors.

#### B. Shape and composition fluctuations

All measurements of the Bik1 surface and bulk fluctuations were taken with a 60x Water Immersion objective (N.A 1.3). We observe shape fluctuations due to low interfacial tension for Bik1 droplets at very low partition coefficients as seen in Fig. S20. As discussed in the main text, we can use bright-field microscopy kymographs to look for sustained density fluctuations in the sample. We observe streaks in the kymograph indicating that the composition fluctuations are somewhat long lasting. Supplemental Movie 4 shows the movie of the fluctuating droplets.

### S5. LIQUID-VAPOR COEXISTENCE OF WATER

All of the data for water is taken from the NIST database [62, 63].

**FIG. S20.**
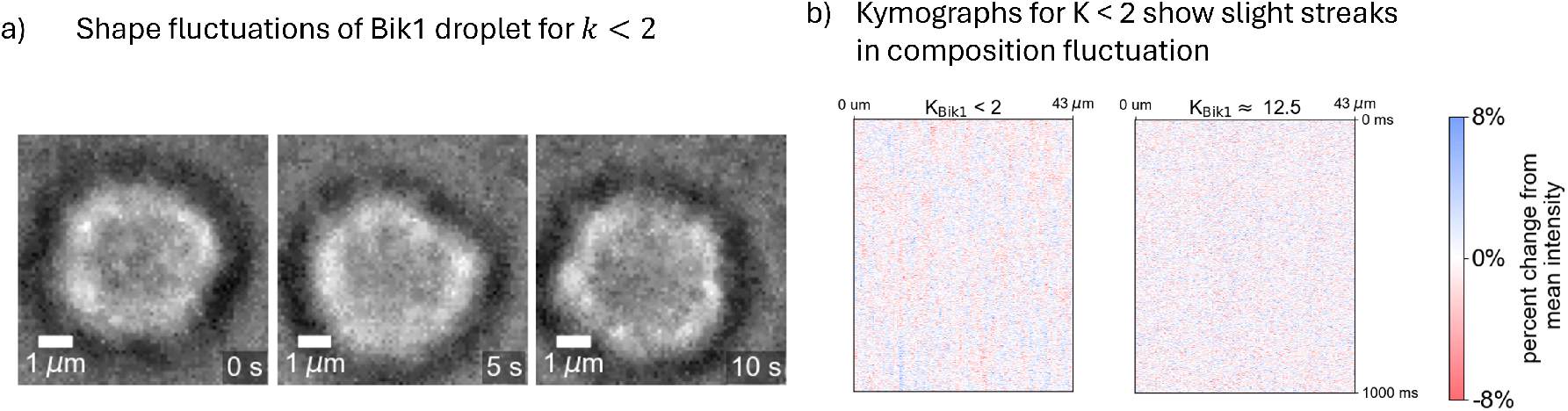
a) Shape fluctuations of Bik1 condensates with partition coefficient smaller than 2 (Composition: 500*µ*M Bik1, 385mM NaCl). We cannot reliably measure the partition coefficient for this composition due to the interaction of the laser with the low-*k* condensates (brightness and contrast adjusted) . b) Kymographs of the dilute phase for two different condensate partition coefficients. We observe qualitatively similar streaks in the low partitioning case though they are not as stark as the BSA/PEG condition.

#### A. Fit to the phase diagram

According to the 3D Ising model phase boundary (also the law of coexisting densities and rectilinear diameters [106]), the dense and dilute phase compositions have the following form:

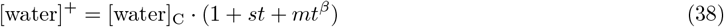

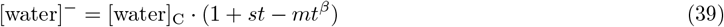

where +*/*− represents the dense/dilute phase and *β* = 0.3264. t is the normalized temperature given by the equation:

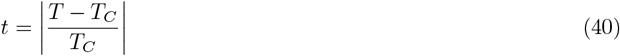

There are four fitting parameters in this equation – {[water]_C_, *s, m, T*_C_ }. We use a least squares fitting algorithm to find the best values of the fit parameters to fit the data.

There is a gauge invariance in the equations 38, 39 which has to do with the overall scale of *t*.

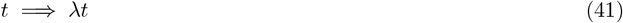

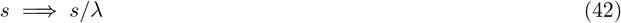

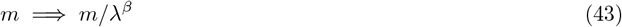

The following transformation leaves the equations 38, 39 invariant.

Assuming the form of *t* (Eq. 40), we have fixed the overall scale of *t* and therefore fixed the gauge.

#### B. Getting Isothermal compression modulus of water from NIST database

In the NIST database [62], the isothermal compression modulus of water is not readily available. It provides the molar density of water *n* (moles/L), speed of sound (*v*_sound_), specific heat capacity at constant volume (*c*_*v*_) and constant pressure (*c*_*v*_) for both the liquid and vapor phases as a function of temperature. From the molar density of water, we compute the mass density of water in both phases by multiplying with the molar mass of water: 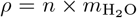 We use this to compute the isothermal compression moduli of the liquid and vapor phases in the following way:

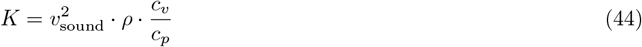

#### C. Scaling of interfacial tension and compressional modulus with normalized temperature

The scaling of interfacial tension and compressional moduli vs *t* is predicted using the 3D Ising model. This is a well known result from classical critical phenomena literature [74].

**FIG. S21.**
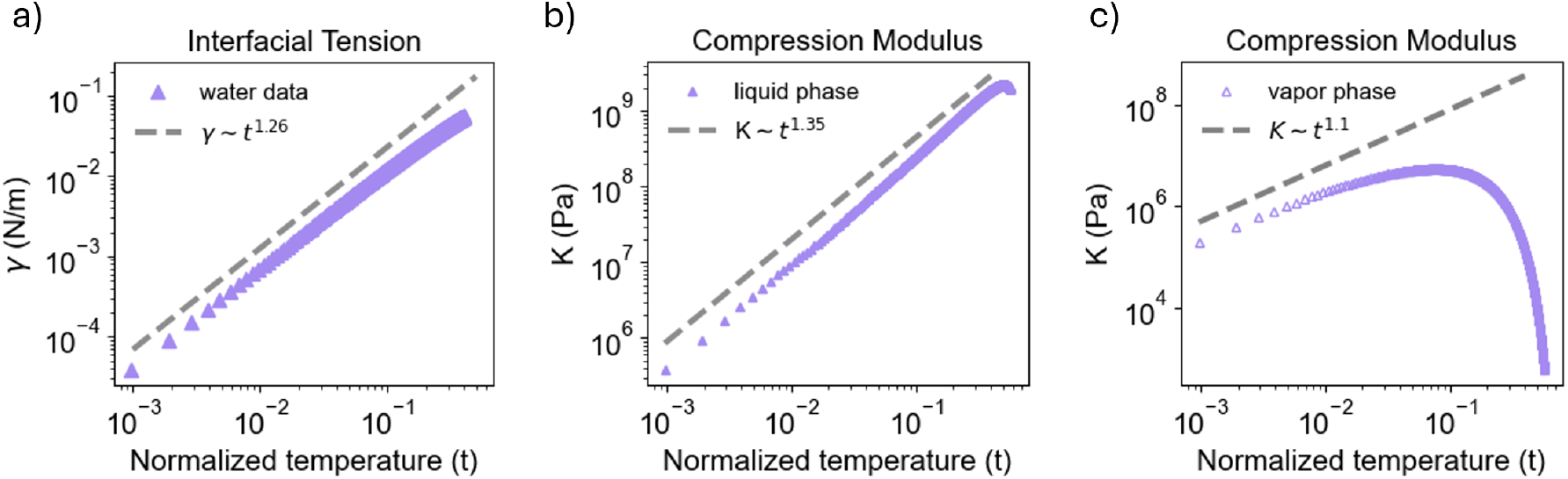
Scaling of material properties of liquid-vapor coexistence of water with normalized temperature. Interfacial tension of liquid-vapor interface as a function of normalized temperature (blue triangles) with the predicted exponent *µ* shown in gray dashed line. b) Compression modulus of the liquid phase along with a fit power law exponent *K* ∼ *t* ^*µ*^ . c) Compression modulus of the vapor phase along with the predicted power law exponent shown in gray dashed line.

### S6. UNIVERSALITY

#### A. Connecting *k* **and** *t* = Θ(*k*)

In this section, we relate the normalized temperature *t* traditionally used in critical phenomena to the partition coefficient *k* via the relation *t* = Θ(*k*) introduced in Fig. 5c), and illustrate in practice that the (experimentally far more convenient) variable *k* is at least as useful in describing both our data (Fig. 5c)) and more traditional critical points (Fig. S23).

We shall argue that using an empirical form appears to work spectacularly well in practice. As support for our empirical form, we start with a somewhat technical calculation of Θ(*k*) valid near the critical point, using the coexistence boundaries predicted by theory (equations 38 and 39). We see that *k* = [*X*]^+^*/*[*X*]^−^ = (1 + *st* + *mt*^*β*^)*/*(1 + *st* − *mt*^*β*^). Taking *β ≈* 1*/*3, this becomes a cubic equation with the general solution For *s* not equal to zero, we could just invert the function numerically. As it happens, if we approximate *β* = 1*/*3, we can cast the problem as the root of a cubic polynomial, and we find the unintuitive solution

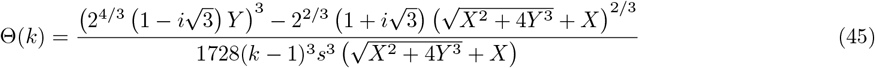

where

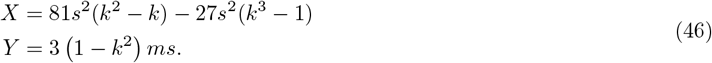

This form can agree well with our data for *t* = Θ(*k*) *<* 0.3 where *k ≈* 80, but by then *X*^−^ has become negative. The slope *s* is only the first of the corrections to scaling expected this far from the critical point. We would need to add the ideal-gas entropy of the dilute phase (repelling from the zero-concentration boundary) to make a more realistic estimate.

This inspires us, in the spirit of fitting universal scaling functions, to simply find the simplest functional form which fits the data.

The empirical form proposed in the main text

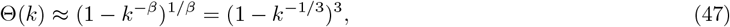

(entirely consistent with the universal properties near the critical point) was designed to fit our experimental data in Fig. 5c). We see in Figure S22 that it also collapses experimental coexistence curve data for a variety of simple liquids, with small deviations for *k >* 100.

**FIG. S22.**
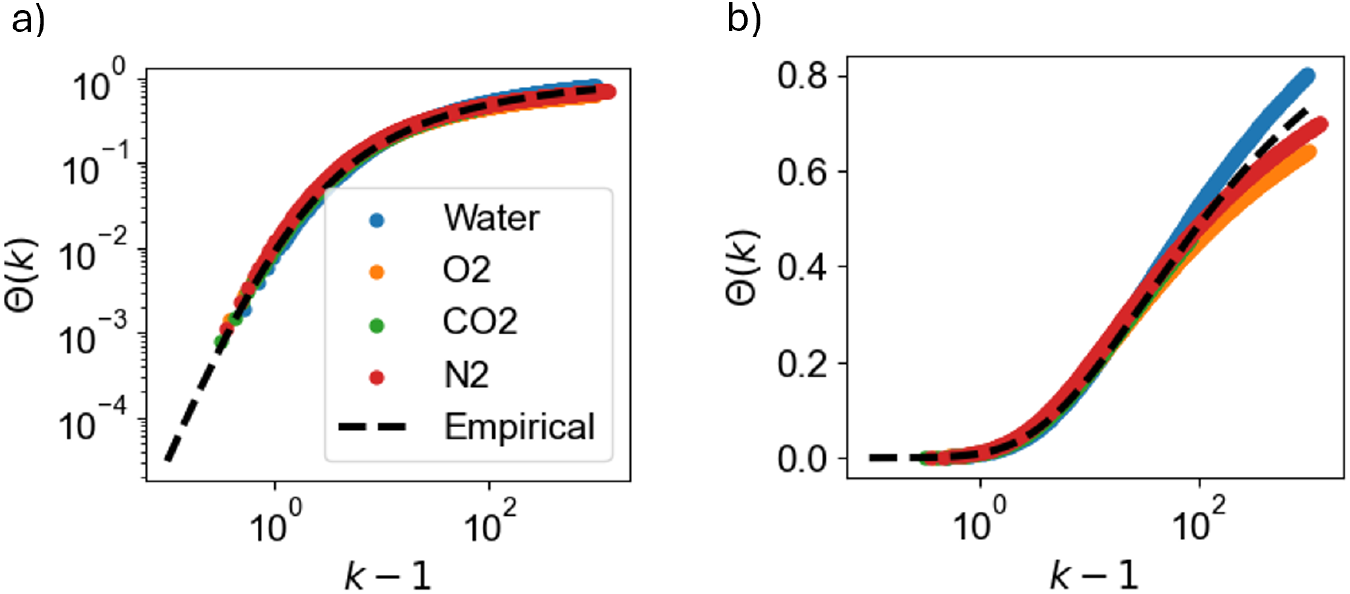
a) Data with empirical form of Θ(*k*) b) Linear-log plot shows that the form deviates after *k* ≈100

#### B. Partition coefficients capture universality as faithfully as normalized temperature

We see in Fig. S23 that we can describe the interfacial tensions and the compression moduli both near and far from the critical point *k* = 1, *t* = 0 (see top and bottom plots of these properties for water, oxygen, nitrogen and carbon dioxide [62], equally well using *t* and *k*. (Using *t* does exhibit the straight-line universal power-law behavior for a longer region than does *k*, but both are equally predictive.)

**FIG. S23.**
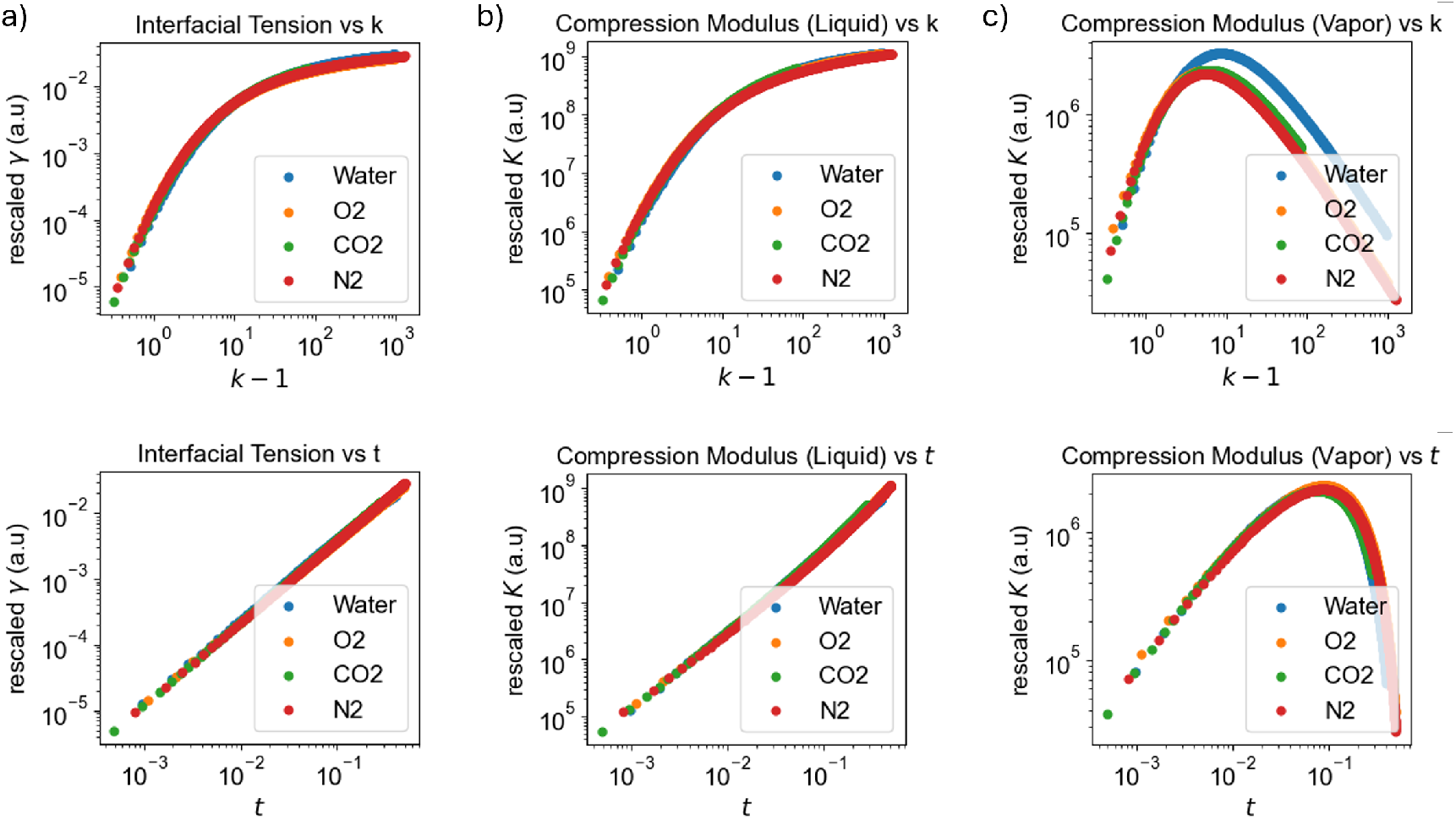
a) Rescaled interfacial tension collapse with partition coefficient *k* (upper) and scaled normalized temperature *t* (lower) b) Rescaled compression modulus of the liquid phase collapse with partition coefficient *k* (upper) and normalized temperature *t* (lower) c) Rescaled compression modulus of the vapor phase collapse with partition coefficient *k* (upper) and normalized temperature *t* (lower).

